# Examining the role of oxygen-binding proteins on the early evolution of multicellularity

**DOI:** 10.1101/2023.12.01.569647

**Authors:** Whitney Wong, Pablo Bravo, Peter J. Yunker, William C. Ratcliff, Anthony J. Burnetti

## Abstract

Oxygen availability is a key factor in the evolution of multicellularity, as larger and more sophisticated organisms often require mechanisms allowing efficient oxygen delivery to their tissues. One such mechanism is the presence of oxygen-binding proteins, such as globins and hemerythrins, which arose in the ancestor of bilaterian animals. Despite their importance, the precise mechanisms by which oxygen-binding proteins influenced the early stages of multicellular evolution under varying environmental oxygen levels are not yet clear. We addressed this knowledge gap by heterologously expressing the oxygen binding proteins myoglobin and myohemerythrin in snowflake yeast, a model system of simple, undifferentiated multicellularity. These proteins increased the depth and rate of oxygen diffusion, increasing the fitness of snowflake yeast growing aerobically. Experiments show that, paradoxically, oxygen-binding proteins confer a greater fitness benefit for larger organisms under high, not low, O_2_ conditions. We show via biophysical modeling that this is because facilitated diffusion is more efficient when oxygen is abundant, transporting a greater quantity of O_2_ which can be used for metabolism. By alleviating anatomical diffusion limitations to oxygen consumption, the evolution of O_2_-binding proteins in the oxygen-rich Neoproterozoic may have been a key breakthrough enabling the evolution of increasingly large, complex multicellular metazoan lineages.

## Introduction

The first unequivocal evidence for macroscopic multicellular animal lineages appears in the fossil record of the Ediacaran, 635 million years ago. Prior to this, the Proterozoic Eon (2.5 billion to 542 million years ago), colloquially known as the "boring billion," bears little trace of complex multicellular life, with only microbial mats and relatively simple multicellular algae inhabiting the oceans[1–7]. It has long been hypothesized that the dramatic rise in atmospheric oxygen levels marking the end of the Proterozoic, from ∼1% of modern atmospheric levels to near-modern concentrations, is linked to the evolution of animal multicellularity[8–10]. With more abundant oxygen, larger organisms would have been expected to be able to overcome diffusion limitations and achieve higher metabolic rates.

Somewhat paradoxically, recent work has shown that increasing oxygen availability can be a powerful force *constraining* the evolution of increased multicellular size. Oxygen is a crucial metabolic cofactor, increasing ATP returns from metabolism and allowing otherwise unfermentable carbon sources to be utilized for growth[11, 12]. However, the evolution of large multicellular bodies creates a strong barrier to diffusion. This can limit the ability of interior cells to access oxygen, reducing their growth rates. The oxygenation of Earth’s atmosphere thus results in counterintuitive evolutionary dynamics, constraining the evolution of macroscopic multicellular size by generating a novel and powerful trade-off between organismal size and growth rate[13, 14] that did not previously exist.

To overcome these anatomical diffusion barriers and support the evolution of large body size, modern animals have evolved specialized oxygen binding and transport proteins such as tetrameric hemoglobins, dimeric hemerythrins, and multimeric hemocyanins. These proteins are all able to bind and release oxygen cooperatively as oxygen levels change as hemoglobin does[15, 16], enabling rapid loading at a respiratory organ and rapid unloading in respiring tissue. These familiar respiratory proteins are components of circulatory systems that thus enable rapid bulk transport of oxygen bound to carriers throughout the body, enabling the evolution of large, active organisms.

However, phylogenetic studies tracing the origins of animal respiratory proteins reveal that freely circulating oxygen carriers with cooperative oxygen binding in blood and hemolymph evolved well after the origin of macroscopic size, and are in fact derived from more ancient monomeric stationary globins and hemerythrins that were not transported in circulatory fluids[17–20]. These more primitive proteins were expressed in body cells and tissues, but still served to facilitate oxygen diffusion from cell to cell and through tissues by increasing the effective diffusion rate of oxygen much as modern tissue globins such as myoglobin do today[21–28]. The ancestral genes encoding these simple intracellular oxygen diffusion facilitators likely originated prior to most recent common ancestor of bilateria[17, 19, 29–31], a critical clade in animal evolution which is defined by the origin of triploblastic embryos with endoderm, ectoderm and mesoderm germ layers which give rise to complex tissues more than two cells thick [32].

The selective drivers favoring the origin of facilitated diffusion via static O_2_-binding proteins remain unresolved. While it seems clear that they provide a benefit by increasing O_2_ diffusion, no prior work has directly examined this benefit as a function of exogenous O_2_ concentration and organism size. We examine these dynamics through synthetic biology and mathematical modeling, heterologously expressing myoglobin from sperm whales (*Physeter macrocephalus*)[33] and myohemerythrin from peanut worms (*Themiste zostericola*)[34] in snowflake yeast (modified *Saccharomyces cerevisiae*), a model system of diffusion-limited multicellularity capable of rapid *in vitro* evolution[35–38]. Our experiments show that, as expected, oxygen-binding proteins significantly enhance oxygen diffusion and total oxygen flux into multicellular yeast clusters. However, as we examine the fitness consequences of facilitated diffusion we find an unexpected result: O_2_-binding proteins confer the greatest advantage for large organisms not when oxygen is rare and most limiting to metabolism, but when it is abundant. Mathematical modeling provides insight into this counterintuitive result, showing that facilitated diffusion leads to greater increase in oxygen flux through the surface of a cluster when oxygen levels are high compared to when they are low, alleviating anatomical diffusion limitations to an abundant resource rather than compensating for low total O_2_ availability. By directly examining the fitness consequences of proteins driving facilitated oxygen diffusion to changes in organismal size and the availability of environmental oxygen, our results provide novel context for the evolution of O_2_-binding proteins in the metazoa.

## Results

To investigate whether heterologously expressed oxygen-binding proteins can enhance oxygen diffusion in our snowflake yeast model system, we first needed to quantify oxygen penetration depth into multicellular clusters. Prior work has shown that internal cells in snowflake yeast are strongly diffusion limited in a low oxygen environment, with only peripheral cells able to respire actively[13].

To examine the effect of O_2_-binding proteins on the depth of oxygen diffusion, we constructed snowflake yeast strains expressing either myoglobin or myohemerythrin integrated at the HO locus (Figure 1A), and introduced the MitoLoc reporter system (preSU9-GFP + preCOX4-mCherry) to visualize aerobic respiration. The MitoLoc system uses a constitutive GFP tag on the outer mitochondrial membrane protein TOM70 to visualize mitochondrial morphology, and colocalization of an mCherry tag on the inner membrane COX4 protein to detect functional membrane potential and respiration[39].

**Figure 1.**
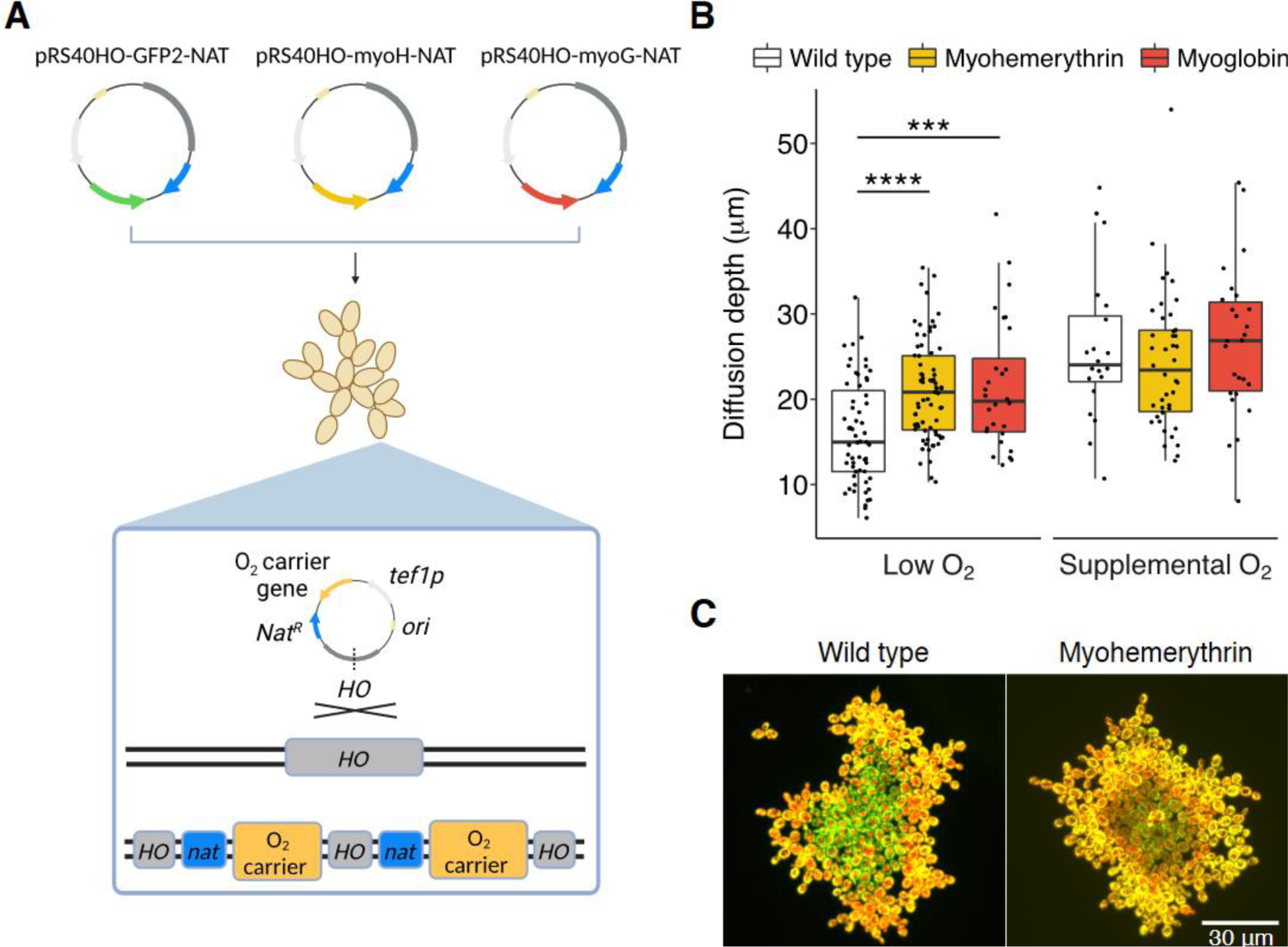
Heterologous expression of oxygen binding proteins increases O_2_ diffusion depth in snowflake yeast. (A) Schematic showing construction of snowflake yeast strains expressing myoglobin or myohemerythrin integrated at the HO locus. (B) Quantification of oxygen diffusion depth using the mitochondrial MitoLoc reporter in snowflake yeast clusters expressing oxygen binding proteins or wildtype controls under oxygen limitation. Myohemerythrin and myoglobin expression significantly increased oxygen diffusion depth, but only in the low oxygen environment. Data are mean ± s.d. (C) Representative fluorescence micrographs of MitoLoc showing increased depth of oxygen penetration (yellow) in clusters expressing myoglobin (right) compared to a wildtype control (left) under low oxygen.

We imaged snowflake yeast clusters expressing the oxygen binding proteins or wildtype controls containing MitoLoc after obligately growth in Yeast Extract Peptone Glycerol media (YEP-Glycerol) under both low oxygen (10 mL cultures shaken without aeration) or supplemental oxygen levels (cultures aerated with room air bubbled through the culture; see Supplemental Figure S1 for typical oxygen traces in each environment). Strains expressing O_2_-binding proteins showed significantly increased depth of oxygen diffusion under oxygen limitation to wildtype controls (Figure 1, 21 µm mean diffusion depth for both myoglobin and myohemerythrin, vs 16 µm for the ancestor; Figure 1B, *p*<0.001, *F*_2,155_=12.57 one-way ANOVA, pairwise comparisons via Tukey’s HSD with α = 0.05), indicating the heterologous proteins enhance penetration of oxygen to internal cells (Figure 1B&C). No significant difference in O_2_ diffusion depth was observed under supplemental oxygen, however, where diffusion limitation is already reduced (*p*=0.48, *F*_2,85_=0.74 one-way ANOVA with α = 0.05).

The fitness effects of expressing oxygen-binding proteins should depend on both the concentration of oxygen in the environment and the size of the multicellular cluster expressing these proteins. Higher environmental oxygen levels should increase depth that oxygen penetrates, while increasing the size of the cluster should reduce the proportion of cells that are oxygenated. We examined this experimentally, by competing myohemerythrin and myoglobin-expressing strains against their ancestors without oxygen-binding protein expression during aerobic growth on YEP-Glycerol. We engineered a small (∼10 μm radius) cluster variant to test how size impacts the fitness advantage of oxygen binding protein expression. Small clusters were generated through deletion of the *BUD8* landmark polarity gene, which gives rise to rounder, smaller snowflake cells, and markedly smaller groups (∼1.8 and ∼5.8 times smaller in radii and volume, Figure 2A and Supplemental Figure S2). Both small and large clusters were competed under the same two oxygen regimes: low (cultures shaken without aeration) and supplemental (cultures aerated with room air bubbled through the liquid). In low oxygen, the majority of the daily culture cycle is spent under 5% saturation at present atmospheric levels. In contrast, under supplemental oxygen (see Methods for details), the vast majority of the daily culture cycle is spent above 50% saturation (Supplemental Figure S1).

**Figure 2.**
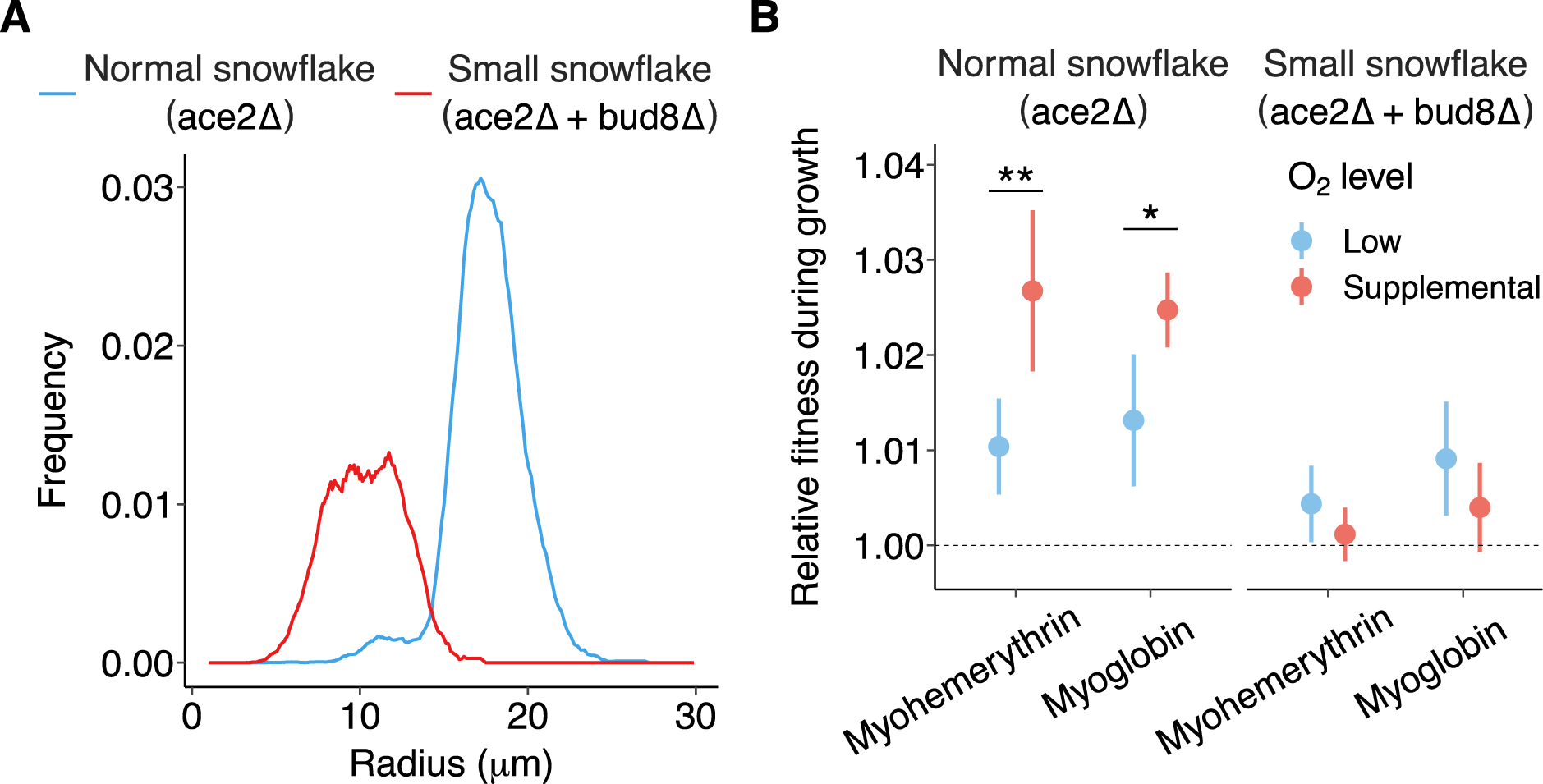
O_2_-binding proteins are most beneficial for large snowflake yeast in a high oxygen environment. (A Small snowflake yeast (*ace2Δ bud8Δ*), engineered by inducing a mutant that causes a random budding pattern to normal snowflake yeast, were approximately half the radius wildtype (*ace2Δ*) snowflake yeast. (B) The benefits of expressing oxygen binding proteins in small-sized snowflake yeast were comparatively modest. Normal-sized snowflake yeast possessed a fitness advantage when expressing O_2_-binding proteins, relative to their isogenic ancestor under both low and supplemental oxygen. For normal sized snowflake yeast, the advantage of expressing O_2_-binding proteins was greater under supplemental oxygen. Dots represent average relative fitness with bars as one standard deviation, *n* = 5 independent competitions for each group.

We calculated the relative fitness of each genotype against a GFP-marked control over three days of growth competition, with 1:100 dilutions carried out daily. In small *bud8Δ* snowflake yeast, myohemerythrin and myoglobin provided no detectable fitness benefit under supplemental oxygen (one-sample t-tests, *p=*0.40, *t=*0.94 and *p=*0.13, *t=*1.91, respectively. Figure 2B.) and a modest, marginally significant fitness advantage in low oxygen (*p=*0.07, *t=*2.44 and *p=*0.03, *t=*3.41, respectively). In contrast, in normal snowflake yeast clusters, both myohemerythrin (one-sample t-tests, *p=*0.0098, *t=*4.62 and *p=*0.002, *t=*7.08, low and supplemental oxygen, respectively) and myoglobin (one-sample t-tests *p=*0.01, *t=*4.24 and *p=*0.0001, *t=*14.06 low and supplemental oxygen, respectively) provided a fitness benefit under all oxygen conditions. This advantage was 2.6-fold (two-sample t-test, *p=*0.008, *t=*3.72, myohemerythrin) and 1.9-fold (two-sample t-test *p=*0.016, *t=*3.25, myoglobin) higher under supplemental oxygen relative to low oxygen conditions.

The finding that large snowflake yeast clusters gained the greatest benefit from oxygen binding protein expression under supplemental oxygen rather than intermediate oxygen was surprising, given we expected oxygen limitation to be most severe under low oxygen conditions[10, 40, 41]. To better understand this counterintuitive result, we modeled the interplay between oxygen availability, cluster size, and globin expression using reaction-diffusion equations in the Julia modeling environment (see supplemental code).

We simulated spherical yeast clusters with radii ranging from 5-70 μm, approximating the range of single cells to large (but not yet macroscopic) snowflake yeast that have undergone significant laboratory evolution for increased size[35–37, 42, 43]. Oxygen diffuses into the cluster from the surrounding environment using diffusion parameters derived from flocculating yeast[44], while being consumed aerobically. Metabolism was modeled using Monod kinetics with parameters extracted from the literature on yeast metabolism[45–48], Myoglobin was added to the simulation at concentrations between from 0 to 0.2 mM, a concentration typical of muscle tissue after correcting for cell packing fractions[48, 49]. Oxygen was allowed to bind and unbind from myoglobin as it diffused, according to measured properties of the protein[22, 50, 51]. Each simulation was initiated with a snowflake yeast with a given radius, myoglobin concentration, and external oxygen concentration, and was allowed to run for 100 simulated seconds to reach equilibrium. We collected data on the final profiles of oxygen concentration across the organism, oxygen metabolism, bound and unbound globin concentration, and average metabolic rates.

This computational model allowed us to predict the profile of oxygen penetration and aerobic respiration for different combinations of cluster size, environmental oxygen, and myoglobin expression. The results provide insight into how oxygen carrier proteins can enhance oxygen flux dependent on multicellular geometry and oxygen availability. The results of this simulation were broadly consistent with experimental observations. A typical cluster in our model (Figure 3A) exhibits a high oxygen concentration at its periphery, which rapidly falls as depth increases. This results in an outer region undergoing high rates of aerobic respiration but a hypoxic core with low or no aerobic respiration, consistent with experimental observations (Figure 1B). Notably, the addition of simulated globin proteins to the model results in a significant increase in the quantity of aerobic respiration and total oxygen within the cluster – oxygen-bound globin reaches equilibrium penetrating deeper below the surface of a cluster than high levels of dissolved oxygen does, contributing to aerobic respiration of inner cells. Depending on the concentration of myoglobin, this dissolved myoglobin-bound oxygen and can represent a large fraction of the total oxygen flux in the system.

**Figure 3.**
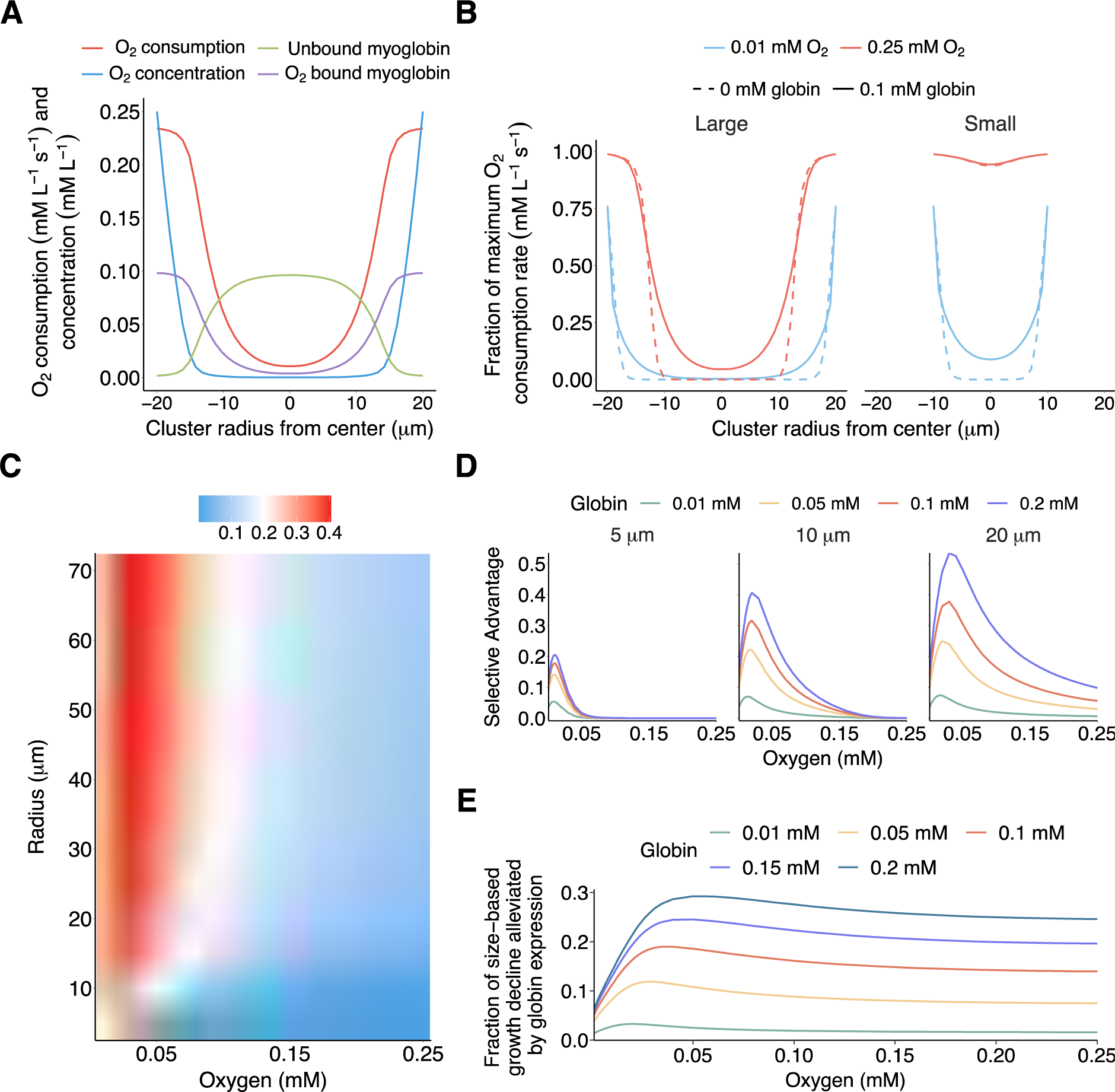
Modeling the relationship between size, environmental oxygen, and the fitness effects of globins. (A) Output of a spherically symmetrical model of the coupled oxygen diffusion, globin diffusion, oxygen/globin binding, and aerobic respiration. Oxygen concentration falls rapidly as distance to the surface of the cluster increases, followed by oxygenated globin concentration, followed by oxygen metabolism. (B) Simulated metabolic rate with depth of large (20 μm radius) and small (10 μm radius) clusters with and without 0.1 mM globin expression. Large clusters attain larger metabolic rate increases in high oxygen environments than low; small clusters are metabolically saturated at high oxygen levels and thus attain no benefit while attaining low benefits at low oxygen levels. (C) The magnitude of globin-induced fitness benefits of clusters at varied radii and oxygen levels at 0.1 mM of globin expression. Fitness benefits are maximized at large radii and intermediate oxygen levels. (d) Graphs of the fitness benefits provided by different degrees of globin expression at 5 μm, 10 μm, and 20 μm radii. At small sizes globin only provides benefits at low oxygen levels with severe diminishing returns to expression; as size increases globin provides larger benefits at intermediate and high oxygen levels without diminishing returns. (E) The fraction of size-induced growth-rate decrease ameliorated by expression of different globin concentrations. As the environmental oxygen level increases, the fraction of size-induced growth retardation that can be reversed by globin expression rapidly increases before leveling off at intermediate values.

Further, the differential effects of oxygen-binding proteins on the growth of clusters of different sizes seen in Figure 2 is captured by this model. Experimentally, we observed that small (∼10 μm radius) clusters exhibited a slight fitness benefit from globin expression at low oxygen levels, but no benefit at high oxygen levels. Our model indicates that at high oxygen levels small clusters are already oxygenated to their core such that globin expression does not appreciably increase the rate of respiration, while at low oxygen levels the expression of myoglobin allows oxygen to penetrate more deeply and the average rate of aerobic respiration to increase (Figure 3B). Our model was similarly conciliant with observations that large (∼20 μm radius) clusters exhibited moderate fitness advantages at low oxygen levels, which rose considerably at higher oxygen levels. Our simulation indicated that large clusters are never oxygenated to their cores (which is consistent with experimental observations, Figure 1B&C) and so always get a fitness benefit from increased diffusion rates, with a larger total benefit arising at higher oxygen levels when facilitated diffusion has a larger impact on overall O_2_ consumption.

Exploring the size and oxygen parameter space revealed an important general trend. Testing all radii from 5 to 70 μm and all oxygen levels from 0.01 to 0.25 mM (Figure 3C, simulated at a myoglobin concentration of 0.1 mM) it is apparent that small clusters never gain a large benefit from expressing oxygen-binding proteins, and the little benefit they do experience is limited to low oxygen levels. As size increases, the maximum benefit obtained from facilitated oxygen diffusion and the oxygen concentration at which that benefit is maximized continually increases. It does, however, plateau at radii large enough that much of the metabolically active cluster is hypoxic. The maximum benefit increases very little above a radius of 40 μm, with the growth rate increase above that size roughly indicating the fold increase in oxygen penetration depth.

Interestingly, we find that for large clusters and high globin concentrations the maximum benefit obtained from globin expression is realized at intermediate oxygen concentrations of approximately 0.05 mM – roughly a fifth of saturation – with smaller but significant benefits remaining at higher oxygen levels. This is unlike the effect in small clusters, in which the benefits of globin expression are maximized at very low oxygen levels and then continually decrease with increasing oxygen (Figure 3D).

Examining the parameter space in more detail, we find that small clusters not only fail to obtain any benefit from globin expression above low oxygen levels, but that at these low levels the benefit that can be obtained from globin expression is limited. As the quantity of globin expressed increases, the benefit obtained rapidly plateaus. However, as cluster size increases, additional globin begins to provide incremental increases to metabolic rate. As size and globin concentrations rise, the oxygen level at which they provide their maximum benefit does as well.

Finally, we examine how much globin can ameliorate the growth costs of multicellularity (that is, the growth of the group relative to that of a single cell). This effect is roughly linearly dependent on the concentration of globin expressed, rising rapidly as environmental oxygen increases from zero before leveling out at intermediate oxygen levels (circa 0.05 mM) and remaining roughly constant above this level (Figure 3E). In our model, expressing 0.25 mM globin can reverse nearly 30% of the cost of increased size.

Taken together, this model reveals three important principles of the effects of globin expression on metabolic rate in respiring multicellular clusters. First, the benefits of globin expression to small clusters are low, and only present at all in low oxygen. Second, the metabolic benefits of globins rise with increasing radius until plateauing when a large enough fraction of the total cluster volume remains hypoxic despite enhanced diffusion. Third, large clusters exhibit continuously increasing incremental benefits from increased globin concentration while small clusters exhibit rapidly diminishing returns. This explains the patterns we observed experimentally (Figure 2) in which expression of oxygen-binding proteins exhibited large fitness benefits in large clusters and small benefits in small clusters with divergent effects of oxygen supplementation, and suggests that facilitated oxygen diffusion provides its largest benefit for large diffusion-limited organisms at high oxygen levels.

## Discussion

Our findings have implications for understanding the role of oxygen-binding proteins as a key innovation facilitating the early evolution of large size and complexity in multicellular organisms. Oxygen availability has long been suspected to be a key factor in the evolution of large complex multicellular organisms, as the great increase in multicellular size and complexity observed after the Neoproterozoic oxygenation event corresponds in time with oxygen rising to near current levels.

Oxygen-binding proteins appear to have originated in the common ancestor of bilaterian animals, a clade characterized by complex anatomy and thick tissues. However, the role of oxygen-binding proteins as a key innovation facilitating the early evolution of large size has remained unclear. Previous studies have suggested that oxygen-binding proteins may have evolved as a response to low oxygen levels in the environment, allowing multicellular organisms to cope with hypoxic stress. However our results challenge this hypothesis, as we found in both experiments and biophysical simulations that oxygen-binding proteins provide relatively little benefit to small organisms, and provide the greatest fitness benefit to larger organisms under higher oxygen levels. This is at first counterintuitive – why does facilitating the diffusion of oxygen not provide the greatest benefit when oxygen is least available? This can be explained by the fact that at high oxygen, any oxygen limitations which exist are *anatomical* rather than *environmental* – while there is copious oxygen available in the environment, diffusion through rapidly metabolizing biomass depletes it before it can reach deeply into tissues.

Our modeling results demonstrate how oxygen binding proteins ameliorate anatomical diffusion limitations. Notably, the addition of globins to a model that previously lacked it (figure 3b) leads to a steepening of the oxygen and aerobic metabolism gradient at the *surface* of a cluster as deoxygenated globins diffuse outwards and take it up, increasing diffusion into the organism from the environment when it is available (see supplemental figure S3 for additional data). The addition of globins also leads to a shallowing of these gradients deeper within the cluster, as bound globins diffuse more deeply and give up their globins to low-oxygen tissues.

We therefore suggest an alternative hypothesis for the role and timing of the origins of oxygen-binding proteins in the animal lineage: that they may have evolved *in response to* rising oxygen levels in the environment, allowing larger multicellular organisms to then exploit the increased metabolic potential of abundant environmental oxygen. Our hypothesis is consistent with the fact that oxygen binding globins and hemerythrins appear to have an origin circa the common ancestor of bilateria[17, 19, 29–31]. These proteins also first originate in the Ediacaran period, after the Neoproterozoic oxygenation event in which oxygen levels rose to near today’s values[40, 52]. An independent origin of oxygen-binding globins appears to have also occurred in land plants with nitrogen-fixing root nodules, where a similarly highly metabolically active non-photosynthetic tissue is also required to maintain high metabolic rates despite diffusion limitations and other oxygen constraints[28, 53].

The advent of dedicated oxygen-binding proteins thus appears tied to a period of rapidly increasing atmospheric oxygen levels, in organisms with thick metabolically active tissues prone to hypoxia from poor oxygen diffusion. These oxygen-binding proteins may have represented a key breakthrough, ameliorating anatomical oxygen limitations inherent to large, dense, metabolically-active multicellular organisms, thereby enabling continuing increases in organismal size and complexity. Our results support the hypothesis that the ancestral emergence of simple intracellular oxygen binding proteins aided early multicellular lineages in overcoming size constraints imposed by diffusion limitation. Rather than evolving as a response to oxygen starvation, oxygen-binding proteins likely emerged because of the oxygen abundance of the Neoproterozoic, allowing early animals to overcome anatomical diffusional limitations and exploit the increased metabolic potential of this abundant resource. By integrating theory and experiments, we this work develops a new perspective on how innovations in oxygen utilization facilitated increases in multicellular scale and complexity, highlighting the importance of global environmental change in shaping the pattern and process of major evolutionary transitions.

## Methods

### Strain construction

All *S. cerevisiae* strains were constructed starting from the homozygous diploid Y55 derivative Y55HD previously used by the Ratcliff laboratory[37] and strain GOB8 bearing an *ACE2* deletion (*ace2Δ::KANMX/ace2Δ::KANMX*) also previously described[13, 36, 37, 43]. We synthesized yeast codon-optimized reading frames coding for peanut worm *(Themiste zostericola*) myohemerythrin[27], sperm whale (*Physeter macrocephalus*)[26] myoglobin, and GFP via ThermoFisher gene synthesis and ligated them into a custom expression vector under the control of a *TEF1* promoter with *NATMX6* resistance cassette and a region of *HO* homology for chromosomal insertion. Plasmids were inserted into the chromosome by cutting within the HO homology using AfeI or BsaAI. This method of integration generates a tandem repeat array of the expression vector integrated into the plasmid at the *HO* locus bracketed by repeats of the *HO* homology region. Due to myoglobin requiring heme to form into a functional protein, we enhanced heme production in myoglobin expression lines and a GFP control line by replacement of the *HEM3* gene promoter (the rate-limiting step in heme production in *S. cerevisiae*) by the *TEF1* promoter and a *HYGMX6* selectable marker via standard PCR-based methods. This generated for ‘normal-sized’ snowflake strains (see strain list). Small-sized yeast for fitness assays were created by first deleting both copies of the *BUD8* gene in snowflake yeast using standard PCR-based methods using the HYGMX6 marker in addition to previously described manipulations, generating four additional strains (see strain list). All genotypes were constructed by standard yeast husbandry techniques involving sporulation and mating of modified strains.

MitoLoc-bearing strains used to visualize and quantify oxygen diffusion were created by inserting a single copy of the *preSU9-GFP* + *preCOX4-mCherry* construct from the MitoLoc plasmid (Addgene #58980) into the same HO-integrating expression vector. This plasmid was cut as previously described for genomic insertion as previously described to generate two heterozygous MitoLoc and oxygen-binding protein expressing snowflake strains (see strain list)

### Plasmids used

pYM25: *GFP* for protein fusion and a *hphNT2* resistance cassette[54]

pWR86: *TEF1* promoter and *hphNT2* for gene overexpression

pWR78: Expression of *P. macrocephalus myoglobin*[33] under a *TEF1* promoter, *NATMX6* resistance, *HO* homology for chromosomal insertion

pWR79: Expression of *T. zostericola myohemerythrin*[34], under a *TEF1* promoter, *NATMX6* resistance, *HO* homology for chromosomal insertion

pWR162: Expression of *GFP* under a *TEF1* promoter, *NATMX6* resistance, *HO* homology for chromosomal insertion

### Oligonucleotides used

Bud8_S2

TACCCAATATCCTCTTTCTACTTGAGAATTTTTTCGATTCTACATGAAGTatcgatgaattcgagctcg

Bud8_R

GACAGAACAGTTTTTTATTTTTTATCCTATTTGATGAATGATACAGTTTCgacatggaggcccagaatac

Hem3_oe_F

AAAGCGAAGAAAATCTAGATAAATTTGTAGTTGGTAAATACACACGTACTccagatctgtttagcttgcc

HEM3_oe_R

ACCGCCAATTTCGATTTTCTCCCACCAATATGTAGAGTTTCAGGGCCCATcttagattagattgctatgctttc

### Strains used

yAB623: Y55HD background, *ace2△:KanMX4/ace2△:KanMX4, ho:MyoH:NatMX6/ho:MyoH:NatMX6*

yAB626: Y55HD background, *ace2△:KanMX4/ace2△:KanMX4, tef1pr-HEM3:HphNT2/tef1pr-HEM3:HphNT2, ho:MyoG:NatMX6/ho:MyoG:NatMX6*

yAB632: Y55HD background, *ace2△:KanMX4/ace2△: KanMX4, tef1pr-HEM3:HphNT2/tef1pr-HEM3:HphNT2, ho:GFP:NatMX6/ho:GFP:NatMX6*

yAB635: Y55HD background, *ace2△:KanMX4/ace2△:KanMX4, ho:GFP:NatMX6/ho:GFP:NatMX6*

yAB714: Y55HD background, *ace2△:Kanmx4/ace2△:Kanmx4, bud8△:HphNT2/bud8△:HphNT2, tef1pr-HEM3: HphNT2/tef1pr-HEM3: HphNT2, ho:MyoG:NatMX6/ho:MyoG:NatMX6*

yAB718: Y55HD background, *ace2△:Kanmx4/ace2△:Kanmx4, bud8△: HphNT2/bud8△:HphNT2, tef1pr-HEM3:HphNT2/tef1pr-HEM3:HphNT2, ho:GFP2:NatMX6/ho:GFP2:NatMX6*

yAB723: Y55HD background, *ace2△:KanMX4/ace2△: KanMX4, bud8△:HphNT2/bud8△:*

*HphNT2, ho:MyoH:NatMX6/ho:MyoH:NatMX6*

yAB727: Y55HD background, *ace2△: KanMX4/ace2△: KanMX4, bud8△: HphNT2/bud8△:*

*HphNT2, ho:GFP2:NatMX6/ho:GFP2:NatMX6*

yAB708: Y55HD background, *ace2△:KANmx4/ace2△:KANmx4, HEM3/tef1pr-HEM3:HphNT2, ho:MyoG:NatMX6/ho:MitoLoc:NatMX6*

yAB710: Y55HD background, ace2△:KanMX4/ace2△:KanMX4, ho:MyoH:NatMX6/ho:MitoLoc:NatMX6

### Imaging MitoLoc

To quantify oxygen diffusion depth, the engineered MitoLoc strains were grown in YEP-Glycerol (1% yeast extract, 2% peptone, 2.5% glycerol) in order to ensure their energy metabolism was fully dependent upon respiration. To vary the amount of available oxygen, 10 mL cultures were grown in a low oxygen environment involving only shaking at 250 RPM at 30°C, and shaking with supplemental oxygen supplied by bubbling room air through the growth media. After 6-7 hours of growth post-inoculation, each strain was imaged using a Nikon Eclipse Ti inverted microscope under TRITC and FITC channels. The diffusion depth of oxygen for each strain was determined by finding the area of cells showing evidence of aerobic respiration due to colocalization between MitoLoc red and green markers using ImageJ.

### Fitness assays

Monocultures of all necessary strains were grown for two days before mixing with a corresponding GFP-expressing control strain at an equal ratio to begin competition assays. Small and large myohemerythrin or myoglobin expressing strains were mixed with an equal volume of small and large GFP-expressing strains and the ratio of clusters of each genotype measured. A total of 100 microliters of this mixture was inoculated into fresh media at the start of each competition. . The populations were grown for a total of three days, inoculating 100 microliters of culture into fresh media every twenty four hours. After measuring the final proportions for each population, we calculated the relative fitness by finding the ratio of Malthusian parameters between the target and control strain[55–57]. The same supplemental oxygen setup from imaging MitoLoc was used to vary environmental oxygen levels. Fluorescence microscopy was used to measure the proportion of the oxygen-binding protein to GFP clusters within a population.

### Mathematical modeling of the dynamics of oxygen, globin, and size

The generic 1-dimensional diffusion equation in cartesian coordinates of the concentration of generic substance *f* with respect to time *t* and position *x* is as follows, where variable *D_f_* corresponds to the diffusion constant of substance *f*, giving us equation 1:

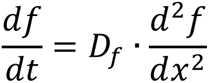

To transform the diffusion equation into 3 dimensional symmetrical spherical coordinates, with *x* now corresponding to a radius from the center of a sphere, we use the following equation 2:

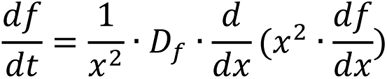

This can be rearranged into the final model’s version of the diffusion equation in polar coordinates, resulting in equation 3.

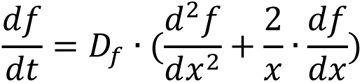

The generic variable *f* can be replaced by the variable *o* for free oxygen concentration, *m_u_* for unbound myoglobin concentration, and *m_b_* for bound myoglobin concentration, all in units of mMol/L. The generic diffusion constant *D_f_* can be replaced by the diffusion constant *D_o_* for the diffusion of oxygen or *D_m_* for the diffusion constant of myoglobin (equal for both bound and unbound to oxygen), both in units of µm^2^s^-1^.

Free oxygen consumption was modeled via the Monod equation[58]. This requires a maximum rate of oxygen consumption *o_max_* in mMol/L/s (noting that the volume given is the volume of modeled multicellular organism including empty space as well as cell volume), and a Monod constant *k_u_* for the concentration of oxygen at which the consumption rate is at half-maximum in mMol/L. Change in oxygen concentration caused by oxygen consumption is given by equation 4.

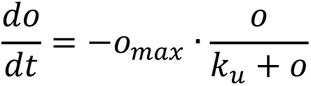

Myoglobin’s interaction with free oxygen is described via an association constant *k_f_* and a dissociation constant *k_r_*. The variable *k_f_* is measured in mM^-1^s^-1^ while the variable *k_r_* is measured in s^-1^, both determined experimentally. The first derivative of free oxygen with time due to association and dissociation with myoglobin is given by equation 5.

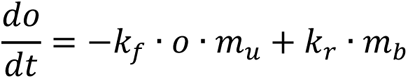

The first derivative of unbound myoglobin with time due to association and dissociation with oxygen is given by equation 6.

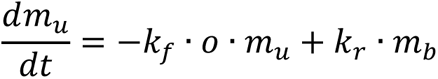

The first derivative of bound myoglobin with time due to association and dissociation with oxygen is given by equation 7.

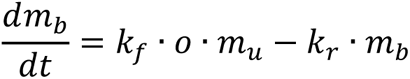

To define the full differential equation for oxygen concentration with respect to time, we add the equations for the first derivatives of oxygen with respect to time as a result of diffusion, association/dissociation, and oxygen consumption, resulting in equation 8.

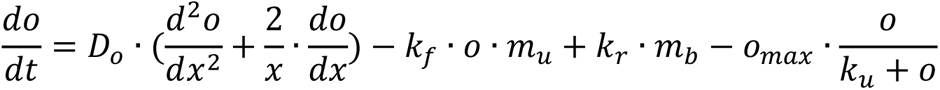

To define the full differential equation for unbound myoglobin concentration with respect to time, we add the equations for the first derivatives of unbound myoglobin with respect to time as a result of diffusion and association/dissociation, resulting in equation 9.

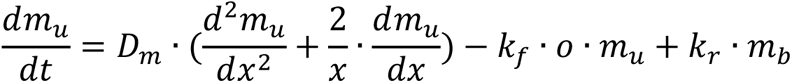

To define the full differential equation for bound myoglobin concentration with respect to time, we add the equations for the first derivatives of bound myoglobin with respect to time as a result of diffusion and association/dissociation, resulting in equation 10.

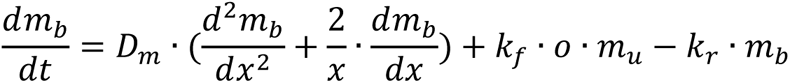

### Parameter values used

While total myoglobin concentration *m_t_* (equal to the sum of *m_b_* and *m_u_*), external oxygen *o_ext_*, and radius *r* were variables reinitialized with every model run, most variables in this model were constants and were obtained from a survey of the scientific literature. Myoglobin association constant *k_f_* and dissociation constant *k_r_* were calculated for heart myoglobin by Endeward, 2012[22] based on numbers from Antonini, 1965[51]. Association constant *k_t_* was taken to be 15400 mM^-1^s^-1^, and dissociation constant *k_r_* was taken to be 60 s^-1^.

While the diffusion constant of oxygen in pure water is measured as over 2000 µm^2^s^-1^[59], diffusion is highly limited in the crowded and heterogenous interior of a cell with many cell walls to cross. Vicente at al, 1998[44] has directly measured the diffusion constant of dissolved oxygen in dense yeast flocs, with variations in experimental technique resulting in measurements between 4.9 and 29.3 µm^2^s^-1^. We used the average of their measurements, 17.1 µm^2^s^-1^. Similarly, the diffusion constant of myoglobin is taken to be 16 µm^2^s^-1^, as measured in the crowded environment of muscle tissue by Papadopoulus et al., 2000[50].

The monod constant *k_u_* of yeast oxygen consumption is provided by model fitting of experimental data by Sonnlietner & Käppelis, 1986[45] as 0.1 mg/L, equivalent to 0.003125 mMol/L. Yeast maximal metabolic rates are seldom measured in terms of the required units of flux per unit cellular volume, however. It is more frequently measured in oxygen flux per unit dry mass. The maximum value of oxygen consumption per unit dry mass of yeast was taken to be 8 mMol/g/hr, also from Sonnlietner & Käppelis, 1986[45]. To convert oxygen consumption rate per unit dry mass of yeast into oxygen consumption rate per unit volume of a multicellular cluster, a dry mass fraction of total mass, a density, and packing fraction is additionally required.

The water mass fraction of yeast is given as 0.604 by Illmer et al, 1999[46], resulting in a dry mass fraction of 0.396. The density of live Y55 yeast is given as 1.1126 g/mL by Baldwin & Kubitschek, 1984[47]. Multiplying these together with the maximum oxygen consumption rate per unit dry mass gives a maximum oxygen consumption rate per unit cellular volume of 0.791 mMol/L/s. However, the snowflake yeast being modeled are not 100% tightly packed cellular volume, instead having a packing fraction resulting from significant free space between cells. According to the dissertation of Dahaj, 2021[48], unevolved snowflake yeast such as those measured in this work have a packing fraction of approximately 0.3. Multiplying this packing fraction by the maximal oxygen consumption rate per unit cellular volume gives a final *o_max_* value of 0.237 mMol/L/s.

### Parameter table

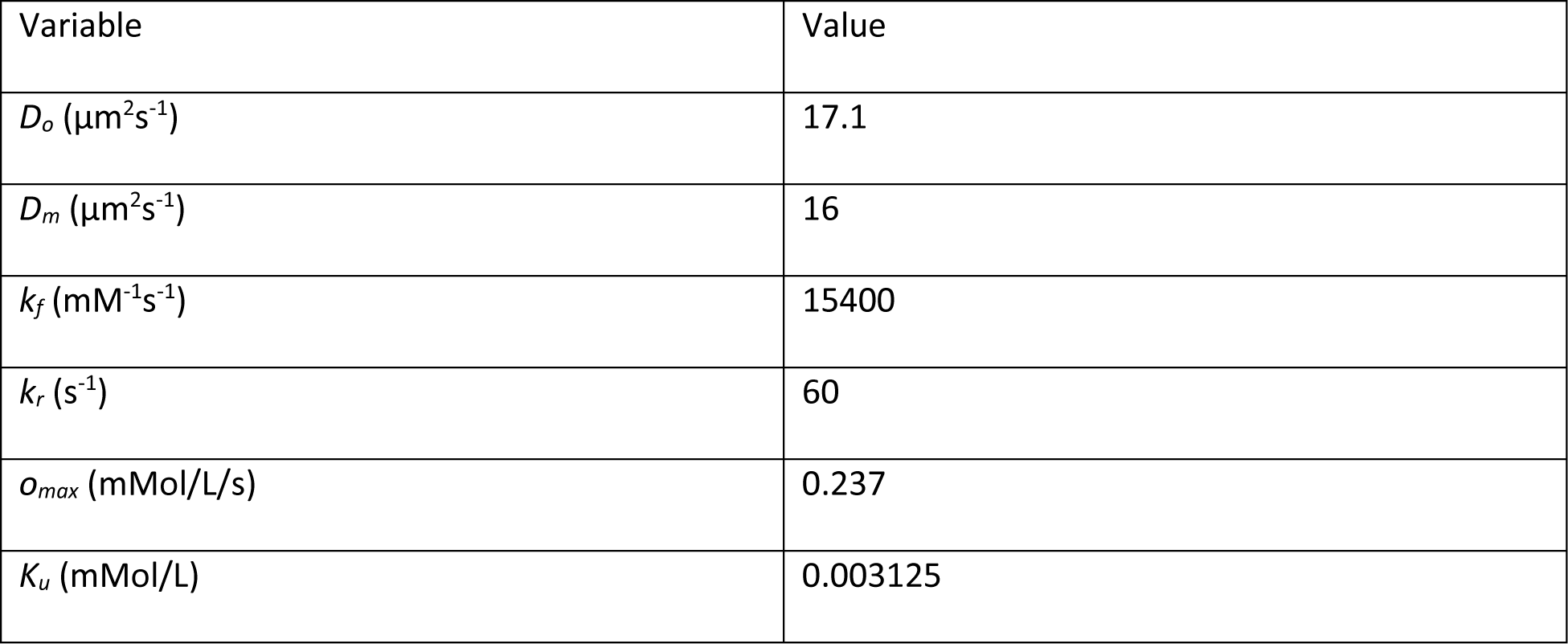

### Numerical simulations

Equations 8-10 were implemented as a set of coupled differential equations in Julia version 1.8.2. The Julia model is provided in the supplemental materials in the form of a Jupyter notebook, project file, and manifest file. In all models the internal oxygen concentration *o* at all points was initialized at 1e-5 mM, and the bound myoglobin concentration *m_b_* was initialized in equilibrium with this. The initial unbound myoglobin concentration *m_u_* at all points was initialized to be equal to the total remaining myoglobin concentration required to reach *m_t_*. Total myoglobin concentration *m_t_*, the external dissolved oxygen boundary condition *o_ext_* (also in mMol/L), and the radius of the cluster *r* (in µm) were all variables provided with the initialization of a given model run. Models were run for 100 seconds of simulated time to reach equilibrium. Models were discretized with a radius step size of 1 μm, and the total oxygen consumption rate according to the Monod equation (equation 4) summed across all discrete radii, to produce a total oxygen consumption rate. This total rate was divided by the volume of the cluster to determine the average metabolic flux per unit volume of a cluster of a given radius in a given environment. The variable *o_ext_* was allowed to vary from a low of 0.01 to a high of 0.25 mMol/L, the approximate maximum solubility of oxygen in water at 30 degrees Celsius[60]. The radius *r* was allowed to vary from 5 µm to 100 µm. The total myoglobin *o_ext_* was allowed to vary from 0 to 0.2 mM/L, a value similar to that which is observed in typical animal heart muscle tissue[49] multiplied by the packing fraction of 0.3[48].

## Supporting information

Myoglobin diffusion model code

## Acknowledgements

We would like to thank members of the Ratcliff and Yunker laboratories for constructive feedback on this paper. W.C.R., A.J.B., and W.W. acknowledge support by grants from the NIH (Grant No. 5R35GM138030), Human Frontiers Science Program grant (RGY0080/2020), NSF Division of Environmental Biology (Grant No. DEB-1845363). P.J.Y. acknowledges funding from the NIH National Institute of General Medical Sciences (grant no. 243 1R35GM138354-01)

---

**Figure S1.**
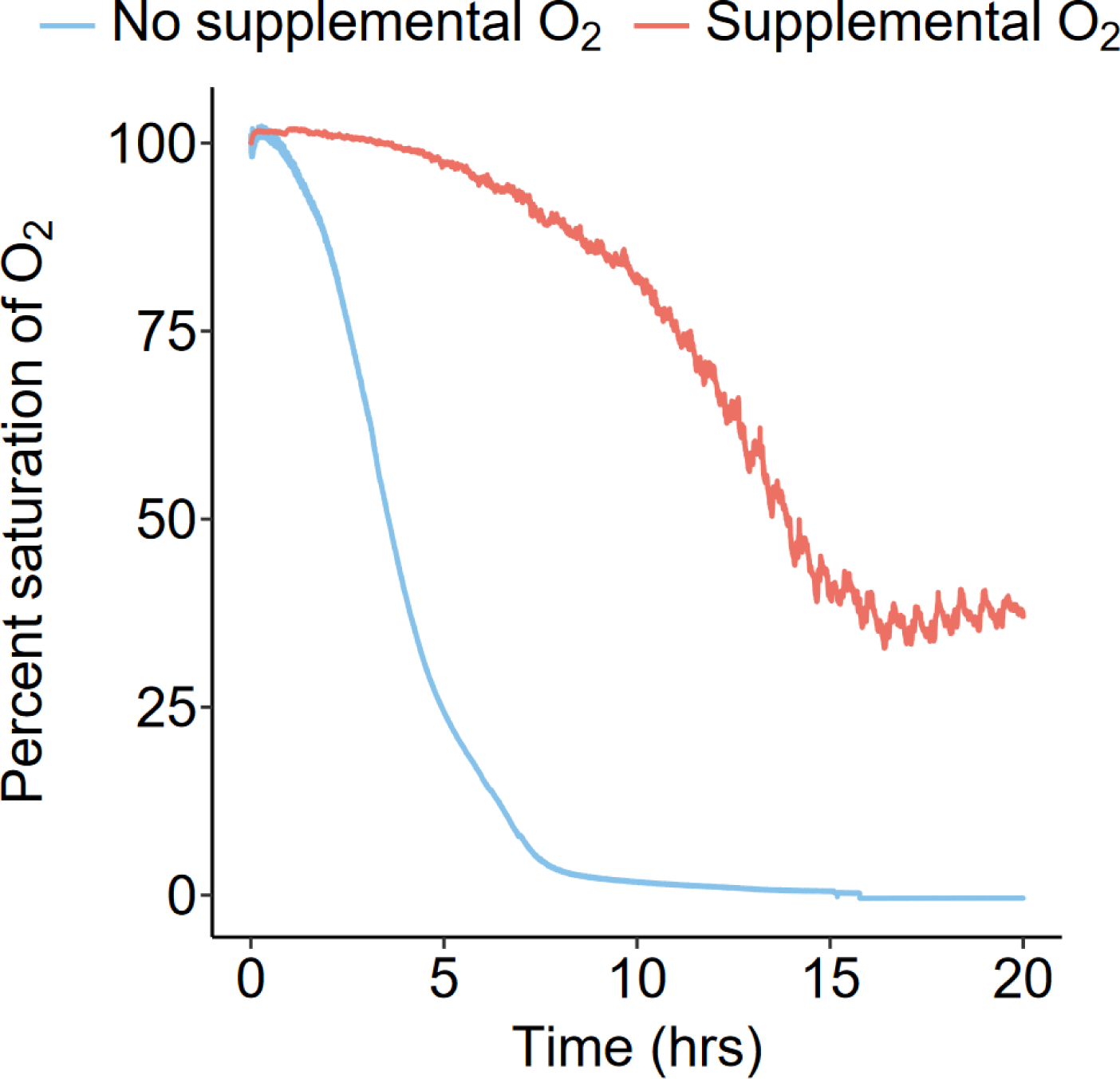
Oxygen levels encountered during ordinary and supplemental oxygen growth cycles. Without supplemental oxygen the majority of a growth cycle is spent at vanishingly low oxygen levels. With supplemental oxygen the majority of a growth cycle is spent at greater than 50% saturation, corresponding to >12.5 mM oxygen. Data averaged from individual runs from Bozdag et al., 2021[13].

**Figure S3.**
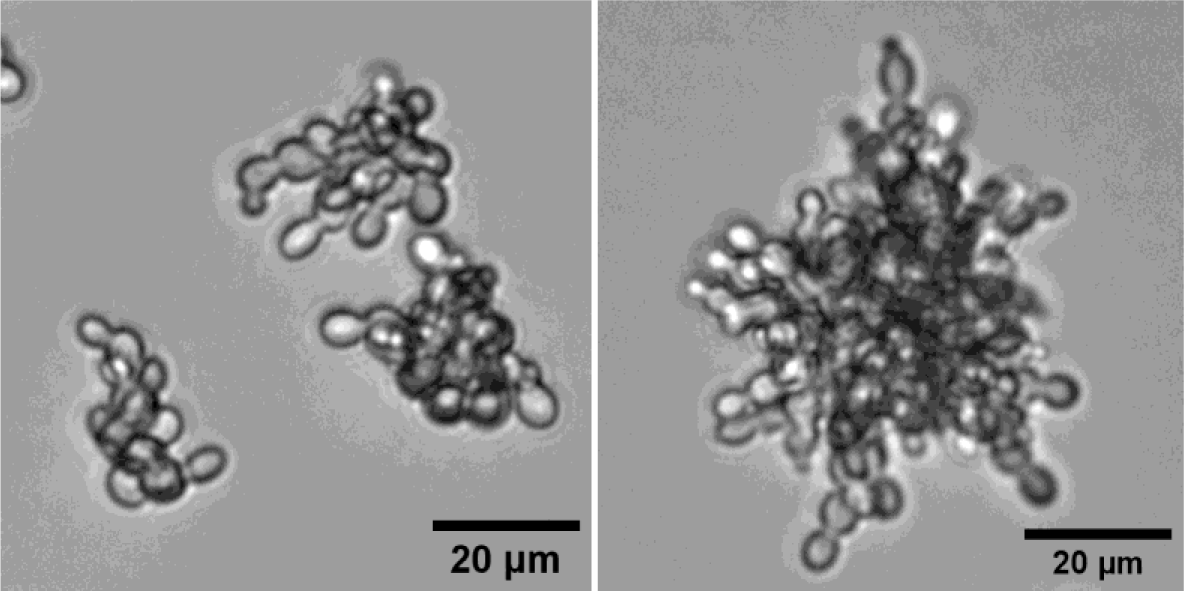
Snowflake yeast (right) and a genetically engineered “Small” strain (bud8Δ, left). Deletion of the *BUD8* gene, which encodes a protein that plays a role in pole selection for bipolar budding, results in daughter cells backbudding towards mother cells thus creating a phenotype of rounder cells for smaller clusters.

**Figure S2.**
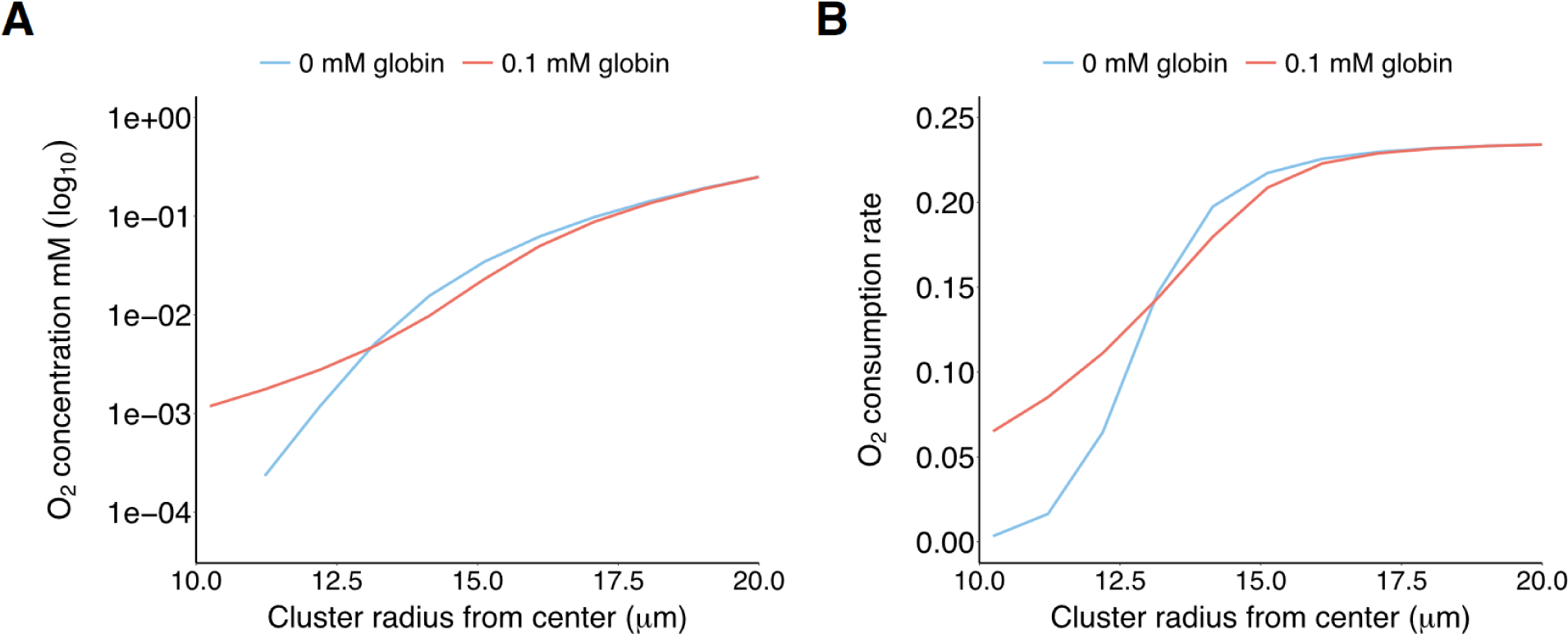
The effect of 0.1 mM globin expression on modeled oxygen gradients within a metabolizing cluster at high (0.25 mM) oxygen concentration. (A) The expression of globin steepens the oxygen gradient at the surface of a cluster, while shallowing the oxygen gradient deeper within a cluster. As the Monod constant of yeast oxygen consumption corresponds to a low oxygen concentration of approximately 3*10^-3^ mM, the oxygen consumption rate (B) of the deeper portions of the cluster with low oxygen is affected much more strongly than the consumption rate of the shallower portions with high oxygen. Thus the increase in oxygen consumption deep within the cluster outweighs the slight decrease in oxygen consumption close to the surface of the cluster.

