## Supplementary material for "Examining the role of oxygen-binding proteins on the early evolution of multicellularity": Myoglobin diffusion model code: 2023-11-10_spherical_parametersweep.ipynb

```
# %% [code]
using ModelingToolkit, MethodOfLines, DifferentialEquations, DomainSets
using Interpolations 
using DataFrames, CSV

# %% [code]
#Make sure you change all diffusion parameter d_oxy!
#water is 2000
#floc is 17.1
#100 is intermediate

# %% [code]
D_oxy = 17.1

# %% [code]
#set the boundaries of the parameter sweep

# %% [code]
#myoglobin
mmin = 0.00
mmax = 0.20
md = 0.01
#oxygen
omin = 0.01
omax = 0.25
od = 0.01
#radius
rmin = 5
rmax = 70
rd = 5

# %% [code]
function integrate_consumption(x_sol, y_sol, radius)
       interp = LinearInterpolation(x_sol, y_sol,extrapolation_bc=Line()) 
       x_integration = Array(0:0.01:radius)
       y_integration = [interp(x) for x in x_integration]
       dx = x_integration[2]-x_integration[1]
       my_int = [y_integration[i]*x_integration[i]*x_integration[i]*dx for i=1:length(x_integration)]
       return sum(my_int)
end

# %% [code]
function run_myo(oxy_outside, myo_in, radius, D_oxy)
       @parameters x t
       @variables o(..) m_u(..) m_b(..) c(..)
       Dt = Differential(t)        # Time derivative
       Dx = Differential(x)        # First spatial der
       Dxx = Differential(x)^2     # Second spatial der
       # This is the set of parameters
       t_min = 0.0                 # Starting point
       t_max = 100                  # s
       #D_oxy = 17.1                # μm^2/s 
       D_m = 16.0                  # μm^2/s  
       k_r = 60.0                  # reverse rate (dissociation constant, 1/s, directly measured)
       k_f = 15.4e3                # forward rate (association constant, 1/mM/s directly measured)
       k_m = k_r/k_f                # km is the reverse rate divided by the forward rate (for prelim values, ~0.004 mM)
       k_u = 0.003125              # O2 uptake Monod constant (mMol/L)
       o_max = 0.237          # Maximum O2 uptake per unit volume (mMol/L/s) (least certain) 
       x_min = -1*radius           # μm derived
       x_max = radius              # μm derived
       #N = 101            # discretization 'bins', ODD NUMBER, dynamically generated!
       N = Int(2*radius+1)
       dx = (x_max-x_min)/N        # bin width

       # Initial conditions
       u0_ext(x, t) = oxy_outside #oxygen
       u0_zero(x, t) = (1e-5) # close to zero
       u0_conc(x, t) = myo_in
       u0_bound(x, t) = oxy_outside/(k_m+oxy_outside)*myo_in
       u0_unbound(x, t) = (1-*(oxy_outside/(k_m+oxy_outside)))*myo_in
       eqs = [Dt(o(x,t)) ~ D_oxy*(Dxx(o(x, t)) + (2/x)*Dx(o(x, t))) - k_f*o(x, t)*m_u(x, t) + k_r*m_b(x, t) - (o_max * o(x, t) / (k_u + o(x, t))),
              Dt(m_u(x, t)) ~ D_m*(Dxx(m_u(x, t)) + (2/x)*Dx(m_u(x, t))) - k_f*o(x, t)*m_u(x, t) + k_r*m_b(x, t),
              Dt(m_b(x, t)) ~ D_m*(Dxx(m_b(x, t)) + (2/x)*Dx(m_b(x, t))) + k_f*o(x, t)*m_u(x, t) - k_r*m_b(x, t)]
       domains = [x ∈ Interval(x_min, x_max),
              t ∈ Interval(t_min, t_max)]
       # Boundary conditions
       bcs = [#o(x, 0) ~ u0_zero(x, 0),    # oxygen = zeros
              #m_u(x, 0) ~ u0_conc(x, 0),  # unbounded = set to initial concentration 
              #m_b(x, 0) ~ u0_zero(x, 0),  # bounded = set to zero
              o(x, 0) ~ u0_ext(x, 0),    # oxygen = external
              m_u(x, 0) ~ u0_unbound(x, 0),  # unbounded = set to initial concentration 
              m_b(x, 0) ~ u0_bound(x, 0),  # bounded = set to initial concentration
              o(x_min, t) ~ oxy_outside,  # At the boundary equal to outside
              o(x_max, t) ~ oxy_outside,  # Same on the other side. 
              Dx(m_u(x_min, t) ) ~ 0,      # For the myoglobins, we set a 'wall'
              Dx(m_u(x_max, t)) ~ 0,      # So the total concentration remains
              Dx(m_b(x_min, t)) ~ 0,      # constant
              Dx(m_b(x_max, t)) ~ 0] 
       @named pdesys = PDESystem(eqs, bcs, domains, [x,t], 
                            [o(x,t), m_u(x,t), m_b(x,t), c(x, t)])
       discretization = MOLFiniteDifference([x=>dx], t, 
                                          approx_order=2, 
                                          grid_align=center_align)
       prob = discretize(pdesys, discretization)
       sol = solve(prob, Rodas5(), save_start=false, save_on=false, save_end=true); # Save only last u 
       #return sol
       consumption = reduce(vcat, o_max .* sol[sol.dvs[2]] ./ (k_u .+ sol[sol.dvs[2]]));
       total_consumption = 4*π*integrate_consumption(Array(sol[x]), consumption, radius)/(radius^3);
       return total_consumption
end

# %% [code]
#setup for with myoglobin
my_sim = run_myo(0.13, 0.01, 5.0, D_oxy)

# %% [code]
Threads.nthreads()

# %% [code]
oxy_list = Array(omin:od:omax)
radius_list = Array(rmin:rd:rmax)
myo_list = Array(mmin:md:mmax)
n_sims = length(oxy_list)*length(radius_list)*length(myo_list)
n_iter_max = Int(ceil((n_sims) / Threads.nthreads()))
sim_iterator = collect(Iterators.product(oxy_list, myo_list, radius_list))
df = DataFrame(CSV.File("output.csv"))
size_current = size(df)[1]
n_iterations = Int(ceil((size_current) / Threads.nthreads()))

# %% [code]
for n in n_iterations:n_iter_max
       O = zeros(Threads.nthreads())
       R = zeros(Threads.nthreads())
       M = zeros(Threads.nthreads())
       C = zeros(Threads.nthreads())
       Threads.@threads for tr in 1:Threads.nthreads()
              i = n*Threads.nthreads() + tr 
              o, m, r = sim_iterator[i]
              c = run_myo(o, m, r, D_oxy)
              O[tr] = o
              R[tr] = r
              M[tr] = m
              C[tr] = c
       end
       tf = DataFrame("Oxygen"=>O, "Radius"=>R, "Myoglobin"=>M, "Consumption"=>C) 
       CSV.write("output.csv", tf,
                 append=true)
end

# %% [code]
```
