## Supplementary material for "Examining the role of oxygen-binding proteins on the early evolution of multicellularity": Myoglobin diffusion model code: Manifest.toml

```
# This file is machine-generated - editing it directly is not advised

julia_version = "1.8.2"
manifest_format = "2.0"
project_hash = "8b80a7d5f7214d696b77d92e8fee525c7da48af5"

[[deps.AbstractAlgebra]]
deps = ["GroupsCore", "InteractiveUtils", "LinearAlgebra", "MacroTools", "Markdown", "Random", "RandomExtensions", "SparseArrays", "Test"]
git-tree-sha1 = "df23d15b1090a3332a09a7a51da45bd9f0a07f92"
uuid = "c3fe647b-3220-5bb0-a1ea-a7954cac585d"
version = "0.27.8"

[[deps.AbstractFFTs]]
deps = ["ChainRulesCore", "LinearAlgebra"]
git-tree-sha1 = "69f7020bd72f069c219b5e8c236c1fa90d2cb409"
uuid = "621f4979-c628-5d54-868e-fcf4e3e8185c"
version = "1.2.1"

[[deps.AbstractTrees]]
git-tree-sha1 = "52b3b436f8f73133d7bc3a6c71ee7ed6ab2ab754"
uuid = "1520ce14-60c1-5f80-bbc7-55ef81b5835c"
version = "0.4.3"

[[deps.Adapt]]
deps = ["LinearAlgebra"]
git-tree-sha1 = "195c5505521008abea5aee4f96930717958eac6f"
uuid = "79e6a3ab-5dfb-504d-930d-738a2a938a0e"
version = "3.4.0"

[[deps.ArgCheck]]
git-tree-sha1 = "a3a402a35a2f7e0b87828ccabbd5ebfbebe356b4"
uuid = "dce04be8-c92d-5529-be00-80e4d2c0e197"
version = "2.3.0"

[[deps.ArgTools]]
uuid = "0dad84c5-d112-42e6-8d28-ef12dabb789f"
version = "1.1.1"

[[deps.ArnoldiMethod]]
deps = ["LinearAlgebra", "Random", "StaticArrays"]
git-tree-sha1 = "62e51b39331de8911e4a7ff6f5aaf38a5f4cc0ae"
uuid = "ec485272-7323-5ecc-a04f-4719b315124d"
version = "0.2.0"

[[deps.ArrayInterface]]
deps = ["ArrayInterfaceCore", "Compat", "IfElse", "LinearAlgebra", "Static"]
git-tree-sha1 = "6d0918cb9c0d3db7fe56bea2bc8638fc4014ac35"
uuid = "4fba245c-0d91-5ea0-9b3e-6abc04ee57a9"
version = "6.0.24"

[[deps.ArrayInterfaceCore]]
deps = ["LinearAlgebra", "SparseArrays", "SuiteSparse"]
git-tree-sha1 = "14c3f84a763848906ac681f94cf469a851601d92"
uuid = "30b0a656-2188-435a-8636-2ec0e6a096e2"
version = "0.1.28"

[[deps.ArrayInterfaceGPUArrays]]
deps = ["Adapt", "ArrayInterfaceCore", "GPUArraysCore", "LinearAlgebra"]
git-tree-sha1 = "fc114f550b93d4c79632c2ada2924635aabfa5ed"
uuid = "6ba088a2-8465-4c0a-af30-387133b534db"
version = "0.2.2"

[[deps.ArrayInterfaceOffsetArrays]]
deps = ["ArrayInterface", "OffsetArrays", "Static"]
git-tree-sha1 = "3d1a9a01976971063b3930d1aed1d9c4af0817f8"
uuid = "015c0d05-e682-4f19-8f0a-679ce4c54826"
version = "0.1.7"

[[deps.ArrayInterfaceStaticArrays]]
deps = ["Adapt", "ArrayInterface", "ArrayInterfaceCore", "ArrayInterfaceStaticArraysCore", "LinearAlgebra", "Static", "StaticArrays"]
git-tree-sha1 = "f12dc65aef03d0a49650b20b2fdaf184928fd886"
uuid = "b0d46f97-bff5-4637-a19a-dd75974142cd"
version = "0.1.5"

[[deps.ArrayInterfaceStaticArraysCore]]
deps = ["Adapt", "ArrayInterfaceCore", "LinearAlgebra", "StaticArraysCore"]
git-tree-sha1 = "93c8ba53d8d26e124a5a8d4ec914c3a16e6a0970"
uuid = "dd5226c6-a4d4-4bc7-8575-46859f9c95b9"
version = "0.1.3"

[[deps.ArrayLayouts]]
deps = ["FillArrays", "LinearAlgebra", "SparseArrays"]
git-tree-sha1 = "4aff5fa660eb95c2e0deb6bcdabe4d9a96bc4667"
uuid = "4c555306-a7a7-4459-81d9-ec55ddd5c99a"
version = "0.8.18"

[[deps.Arrow]]
deps = ["ArrowTypes", "BitIntegers", "CodecLz4", "CodecZstd", "DataAPI", "Dates", "LoggingExtras", "Mmap", "PooledArrays", "SentinelArrays", "Tables", "TimeZones", "UUIDs", "WorkerUtilities"]
git-tree-sha1 = "e97bdb5e241bb57f628968fd56efd9590078ada4"
uuid = "69666777-d1a9-59fb-9406-91d4454c9d45"
version = "2.4.2"

[[deps.ArrowTypes]]
deps = ["UUIDs"]
git-tree-sha1 = "563d60f89fcb730668bd568ba3e752ee71dde023"
uuid = "31f734f8-188a-4ce0-8406-c8a06bd891cd"
version = "2.0.2"

[[deps.Artifacts]]
uuid = "56f22d72-fd6d-98f1-02f0-08ddc0907c33"

[[deps.Atomix]]
deps = ["UnsafeAtomics"]
git-tree-sha1 = "c06a868224ecba914baa6942988e2f2aade419be"
uuid = "a9b6321e-bd34-4604-b9c9-b65b8de01458"
version = "0.1.0"

[[deps.AutoHashEquals]]
git-tree-sha1 = "45bb6705d93be619b81451bb2006b7ee5d4e4453"
uuid = "15f4f7f2-30c1-5605-9d31-71845cf9641f"
version = "0.2.0"

[[deps.AxisAlgorithms]]
deps = ["LinearAlgebra", "Random", "SparseArrays", "WoodburyMatrices"]
git-tree-sha1 = "66771c8d21c8ff5e3a93379480a2307ac36863f7"
uuid = "13072b0f-2c55-5437-9ae7-d433b7a33950"
version = "1.0.1"

[[deps.BFloat16s]]
deps = ["LinearAlgebra", "Printf", "Random", "Test"]
git-tree-sha1 = "a598ecb0d717092b5539dbbe890c98bac842b072"
uuid = "ab4f0b2a-ad5b-11e8-123f-65d77653426b"
version = "0.2.0"

[[deps.BandedMatrices]]
deps = ["ArrayLayouts", "FillArrays", "LinearAlgebra", "SparseArrays"]
git-tree-sha1 = "134fe629f08ca56f07b1d8028736dab437df34f5"
uuid = "aae01518-5342-5314-be14-df237901396f"
version = "0.17.9"

[[deps.BangBang]]
deps = ["Compat", "ConstructionBase", "Future", "InitialValues", "LinearAlgebra", "Requires", "Setfield", "Tables", "ZygoteRules"]
git-tree-sha1 = "7fe6d92c4f281cf4ca6f2fba0ce7b299742da7ca"
uuid = "198e06fe-97b7-11e9-32a5-e1d131e6ad66"
version = "0.3.37"

[[deps.Base64]]
uuid = "2a0f44e3-6c83-55bd-87e4-b1978d98bd5f"

[[deps.Baselet]]
git-tree-sha1 = "aebf55e6d7795e02ca500a689d326ac979aaf89e"
uuid = "9718e550-a3fa-408a-8086-8db961cd8217"
version = "0.1.1"

[[deps.BenchmarkTools]]
deps = ["JSON", "Logging", "Printf", "Profile", "Statistics", "UUIDs"]
git-tree-sha1 = "d9a9701b899b30332bbcb3e1679c41cce81fb0e8"
uuid = "6e4b80f9-dd63-53aa-95a3-0cdb28fa8baf"
version = "1.3.2"

[[deps.Bijections]]
git-tree-sha1 = "fe4f8c5ee7f76f2198d5c2a06d3961c249cce7bd"
uuid = "e2ed5e7c-b2de-5872-ae92-c73ca462fb04"
version = "0.1.4"

[[deps.BitIntegers]]
deps = ["Random"]
git-tree-sha1 = "fc54d5837033a170f3bad307f993e156eefc345f"
uuid = "c3b6d118-76ef-56ca-8cc7-ebb389d030a1"
version = "0.2.7"

[[deps.BitTwiddlingConvenienceFunctions]]
deps = ["Static"]
git-tree-sha1 = "0c5f81f47bbbcf4aea7b2959135713459170798b"
uuid = "62783981-4cbd-42fc-bca8-16325de8dc4b"
version = "0.1.5"

[[deps.BoundaryValueDiffEq]]
deps = ["BandedMatrices", "DiffEqBase", "FiniteDiff", "ForwardDiff", "LinearAlgebra", "NLsolve", "Reexport", "SciMLBase", "SparseArrays"]
git-tree-sha1 = "ed8e837bfb3d1e3157022c9636ec1c722b637318"
uuid = "764a87c0-6b3e-53db-9096-fe964310641d"
version = "2.11.0"

[[deps.CEnum]]
git-tree-sha1 = "eb4cb44a499229b3b8426dcfb5dd85333951ff90"
uuid = "fa961155-64e5-5f13-b03f-caf6b980ea82"
version = "0.4.2"

[[deps.CPUSummary]]
deps = ["CpuId", "IfElse", "Static"]
git-tree-sha1 = "5b735f654bdfd7b6c18c49f1d3ebff34b4b8af43"
uuid = "2a0fbf3d-bb9c-48f3-b0a9-814d99fd7ab9"
version = "0.2.1"

[[deps.CSTParser]]
deps = ["Tokenize"]
git-tree-sha1 = "3ddd48d200eb8ddf9cb3e0189fc059fd49b97c1f"
uuid = "00ebfdb7-1f24-5e51-bd34-a7502290713f"
version = "3.3.6"

[[deps.CSV]]
deps = ["CodecZlib", "Dates", "FilePathsBase", "InlineStrings", "Mmap", "Parsers", "PooledArrays", "SentinelArrays", "SnoopPrecompile", "Tables", "Unicode", "WeakRefStrings", "WorkerUtilities"]
git-tree-sha1 = "c700cce799b51c9045473de751e9319bdd1c6e94"
uuid = "336ed68f-0bac-5ca0-87d4-7b16caf5d00b"
version = "0.10.9"

[[deps.CUDA]]
deps = ["AbstractFFTs", "Adapt", "BFloat16s", "CEnum", "CompilerSupportLibraries_jll", "ExprTools", "GPUArrays", "GPUCompiler", "LLVM", "LazyArtifacts", "Libdl", "LinearAlgebra", "Logging", "Printf", "Random", "Random123", "RandomNumbers", "Reexport", "Requires", "SparseArrays", "SpecialFunctions", "TimerOutputs"]
git-tree-sha1 = "a56dff7bc49b5d5ac43d2c10eb2aef94becd5251"
uuid = "052768ef-5323-5732-b1bb-66c8b64840ba"
version = "3.12.1"

[[deps.CUDAKernels]]
deps = ["Adapt", "CUDA", "KernelAbstractions", "StaticArrays", "UnsafeAtomicsLLVM"]
git-tree-sha1 = "bbab4d1a4001ec322c384dfff0889cec4118da93"
uuid = "72cfdca4-0801-4ab0-bf6a-d52aa10adc57"
version = "0.4.3"

[[deps.Calculus]]
deps = ["LinearAlgebra"]
git-tree-sha1 = "f641eb0a4f00c343bbc32346e1217b86f3ce9dad"
uuid = "49dc2e85-a5d0-5ad3-a950-438e2897f1b9"
version = "0.5.1"

[[deps.ChainRulesCore]]
deps = ["Compat", "LinearAlgebra", "SparseArrays"]
git-tree-sha1 = "e7ff6cadf743c098e08fca25c91103ee4303c9bb"
uuid = "d360d2e6-b24c-11e9-a2a3-2a2ae2dbcce4"
version = "1.15.6"

[[deps.ChangesOfVariables]]
deps = ["ChainRulesCore", "LinearAlgebra", "Test"]
git-tree-sha1 = "38f7a08f19d8810338d4f5085211c7dfa5d5bdd8"
uuid = "9e997f8a-9a97-42d5-a9f1-ce6bfc15e2c0"
version = "0.1.4"

[[deps.CloseOpenIntervals]]
deps = ["ArrayInterface", "Static"]
git-tree-sha1 = "d61300b9895f129f4bd684b2aff97cf319b6c493"
uuid = "fb6a15b2-703c-40df-9091-08a04967cfa9"
version = "0.1.11"

[[deps.CodecLz4]]
deps = ["Lz4_jll", "TranscodingStreams"]
git-tree-sha1 = "59fe0cb37784288d6b9f1baebddbf75457395d40"
uuid = "5ba52731-8f18-5e0d-9241-30f10d1ec561"
version = "0.4.0"

[[deps.CodecZlib]]
deps = ["TranscodingStreams", "Zlib_jll"]
git-tree-sha1 = "ded953804d019afa9a3f98981d99b33e3db7b6da"
uuid = "944b1d66-785c-5afd-91f1-9de20f533193"
version = "0.7.0"

[[deps.CodecZstd]]
deps = ["CEnum", "TranscodingStreams", "Zstd_jll"]
git-tree-sha1 = "849470b337d0fa8449c21061de922386f32949d9"
uuid = "6b39b394-51ab-5f42-8807-6242bab2b4c2"
version = "0.7.2"

[[deps.ColorSchemes]]
deps = ["ColorTypes", "ColorVectorSpace", "Colors", "FixedPointNumbers", "PrecompileTools", "Random"]
git-tree-sha1 = "67c1f244b991cad9b0aa4b7540fb758c2488b129"
uuid = "35d6a980-a343-548e-a6ea-1d62b119f2f4"
version = "3.24.0"

[[deps.ColorTypes]]
deps = ["FixedPointNumbers", "Random"]
git-tree-sha1 = "eb7f0f8307f71fac7c606984ea5fb2817275d6e4"
uuid = "3da002f7-5984-5a60-b8a6-cbb66c0b333f"
version = "0.11.4"

[[deps.ColorVectorSpace]]
deps = ["ColorTypes", "FixedPointNumbers", "LinearAlgebra", "Requires", "Statistics", "TensorCore"]
git-tree-sha1 = "a1f44953f2382ebb937d60dafbe2deea4bd23249"
uuid = "c3611d14-8923-5661-9e6a-0046d554d3a4"
version = "0.10.0"

[[deps.Colors]]
deps = ["ColorTypes", "FixedPointNumbers", "Reexport"]
git-tree-sha1 = "fc08e5930ee9a4e03f84bfb5211cb54e7769758a"
uuid = "5ae59095-9a9b-59fe-a467-6f913c188581"
version = "0.12.10"

[[deps.Combinatorics]]
git-tree-sha1 = "08c8b6831dc00bfea825826be0bc8336fc369860"
uuid = "861a8166-3701-5b0c-9a16-15d98fcdc6aa"
version = "1.0.2"

[[deps.CommonMark]]
deps = ["Crayons", "JSON", "URIs"]
git-tree-sha1 = "86cce6fd164c26bad346cc51ca736e692c9f553c"
uuid = "a80b9123-70ca-4bc0-993e-6e3bcb318db6"
version = "0.8.7"

[[deps.CommonSolve]]
git-tree-sha1 = "9441451ee712d1aec22edad62db1a9af3dc8d852"
uuid = "38540f10-b2f7-11e9-35d8-d573e4eb0ff2"
version = "0.2.3"

[[deps.CommonSubexpressions]]
deps = ["MacroTools", "Test"]
git-tree-sha1 = "7b8a93dba8af7e3b42fecabf646260105ac373f7"
uuid = "bbf7d656-a473-5ed7-a52c-81e309532950"
version = "0.3.0"

[[deps.Compat]]
deps = ["Dates", "LinearAlgebra", "UUIDs"]
git-tree-sha1 = "00a2cccc7f098ff3b66806862d275ca3db9e6e5a"
uuid = "34da2185-b29b-5c13-b0c7-acf172513d20"
version = "4.5.0"

[[deps.CompilerSupportLibraries_jll]]
deps = ["Artifacts", "Libdl"]
uuid = "e66e0078-7015-5450-92f7-15fbd957f2ae"
version = "0.5.2+0"

[[deps.CompositeTypes]]
git-tree-sha1 = "02d2316b7ffceff992f3096ae48c7829a8aa0638"
uuid = "b152e2b5-7a66-4b01-a709-34e65c35f657"
version = "0.1.3"

[[deps.CompositionsBase]]
git-tree-sha1 = "455419f7e328a1a2493cabc6428d79e951349769"
uuid = "a33af91c-f02d-484b-be07-31d278c5ca2b"
version = "0.1.1"

[[deps.Conda]]
deps = ["Downloads", "JSON", "VersionParsing"]
git-tree-sha1 = "6e47d11ea2776bc5627421d59cdcc1296c058071"
uuid = "8f4d0f93-b110-5947-807f-2305c1781a2d"
version = "1.7.0"

[[deps.ConstructionBase]]
deps = ["LinearAlgebra"]
git-tree-sha1 = "fb21ddd70a051d882a1686a5a550990bbe371a95"
uuid = "187b0558-2788-49d3-abe0-74a17ed4e7c9"
version = "1.4.1"

[[deps.CpuId]]
deps = ["Markdown"]
git-tree-sha1 = "fcbb72b032692610bfbdb15018ac16a36cf2e406"
uuid = "adafc99b-e345-5852-983c-f28acb93d879"
version = "0.3.1"

[[deps.Crayons]]
git-tree-sha1 = "249fe38abf76d48563e2f4556bebd215aa317e15"
uuid = "a8cc5b0e-0ffa-5ad4-8c14-923d3ee1735f"
version = "4.1.1"

[[deps.DataAPI]]
git-tree-sha1 = "e8119c1a33d267e16108be441a287a6981ba1630"
uuid = "9a962f9c-6df0-11e9-0e5d-c546b8b5ee8a"
version = "1.14.0"

[[deps.DataFrames]]
deps = ["Compat", "DataAPI", "Future", "InvertedIndices", "IteratorInterfaceExtensions", "LinearAlgebra", "Markdown", "Missings", "PooledArrays", "PrettyTables", "Printf", "REPL", "Random", "Reexport", "SnoopPrecompile", "SortingAlgorithms", "Statistics", "TableTraits", "Tables", "Unicode"]
git-tree-sha1 = "d4f69885afa5e6149d0cab3818491565cf41446d"
uuid = "a93c6f00-e57d-5684-b7b6-d8193f3e46c0"
version = "1.4.4"

[[deps.DataStructures]]
deps = ["Compat", "InteractiveUtils", "OrderedCollections"]
git-tree-sha1 = "d1fff3a548102f48987a52a2e0d114fa97d730f0"
uuid = "864edb3b-99cc-5e75-8d2d-829cb0a9cfe8"
version = "0.18.13"

[[deps.DataValueInterfaces]]
git-tree-sha1 = "bfc1187b79289637fa0ef6d4436ebdfe6905cbd6"
uuid = "e2d170a0-9d28-54be-80f0-106bbe20a464"
version = "1.0.0"

[[deps.Dates]]
deps = ["Printf"]
uuid = "ade2ca70-3891-5945-98fb-dc099432e06a"

[[deps.DecFP]]
deps = ["DecFP_jll", "Printf", "Random", "SpecialFunctions"]
git-tree-sha1 = "a8269e0a6af8c9d9ae95d15dcfa5628285980cbb"
uuid = "55939f99-70c6-5e9b-8bb0-5071ed7d61fd"
version = "1.3.1"

[[deps.DecFP_jll]]
deps = ["Artifacts", "JLLWrappers", "Libdl", "Pkg"]
git-tree-sha1 = "e9a8da19f847bbfed4076071f6fef8665a30d9e5"
uuid = "47200ebd-12ce-5be5-abb7-8e082af23329"
version = "2.0.3+1"

[[deps.DefineSingletons]]
git-tree-sha1 = "0fba8b706d0178b4dc7fd44a96a92382c9065c2c"
uuid = "244e2a9f-e319-4986-a169-4d1fe445cd52"
version = "0.1.2"

[[deps.DelayDiffEq]]
deps = ["ArrayInterface", "DataStructures", "DiffEqBase", "LinearAlgebra", "Logging", "OrdinaryDiffEq", "Printf", "RecursiveArrayTools", "Reexport", "SciMLBase", "SimpleNonlinearSolve", "UnPack"]
git-tree-sha1 = "478408ad9195fce93f2837a18519fbe7f7795b6d"
uuid = "bcd4f6db-9728-5f36-b5f7-82caef46ccdb"
version = "5.40.5"

[[deps.DensityInterface]]
deps = ["InverseFunctions", "Test"]
git-tree-sha1 = "80c3e8639e3353e5d2912fb3a1916b8455e2494b"
uuid = "b429d917-457f-4dbc-8f4c-0cc954292b1d"
version = "0.4.0"

[[deps.DiffEqBase]]
deps = ["ArrayInterfaceCore", "ChainRulesCore", "DataStructures", "Distributions", "DocStringExtensions", "FastBroadcast", "ForwardDiff", "FunctionWrappers", "FunctionWrappersWrappers", "LinearAlgebra", "Logging", "MuladdMacro", "Parameters", "PreallocationTools", "Printf", "RecursiveArrayTools", "Reexport", "Requires", "SciMLBase", "Setfield", "SimpleNonlinearSolve", "SparseArrays", "Static", "StaticArrays", "Statistics", "Tricks", "ZygoteRules"]
git-tree-sha1 = "b410f0b8a52752e1c1723b4316382203f914672c"
uuid = "2b5f629d-d688-5b77-993f-72d75c75574e"
version = "6.113.1"

[[deps.DiffEqCallbacks]]
deps = ["DataStructures", "DiffEqBase", "ForwardDiff", "LinearAlgebra", "Markdown", "NLsolve", "Parameters", "RecipesBase", "RecursiveArrayTools", "SciMLBase", "StaticArrays"]
git-tree-sha1 = "485503846a90b59f3b79b39c2d818496bf50d197"
uuid = "459566f4-90b8-5000-8ac3-15dfb0a30def"
version = "2.24.3"

[[deps.DiffEqGPU]]
deps = ["Adapt", "ArrayInterfaceGPUArrays", "CUDA", "CUDAKernels", "ChainRulesCore", "DiffEqBase", "Distributed", "DocStringExtensions", "ForwardDiff", "KernelAbstractions", "LinearAlgebra", "LinearSolve", "MuladdMacro", "Parameters", "Random", "RecursiveArrayTools", "SciMLBase", "SimpleDiffEq", "StaticArrays", "ZygoteRules"]
git-tree-sha1 = "92f81c7f059160e3f8c2b7d7163532e0eddb67b0"
uuid = "071ae1c0-96b5-11e9-1965-c90190d839ea"
version = "1.25.0"

[[deps.DiffEqNoiseProcess]]
deps = ["DiffEqBase", "Distributions", "GPUArraysCore", "LinearAlgebra", "Markdown", "Optim", "PoissonRandom", "QuadGK", "Random", "Random123", "RandomNumbers", "RecipesBase", "RecursiveArrayTools", "ResettableStacks", "SciMLBase", "StaticArrays", "Statistics"]
git-tree-sha1 = "1a5c145ea1915b92e0d446c05e375f9c69c0348d"
uuid = "77a26b50-5914-5dd7-bc55-306e6241c503"
version = "5.15.2"

[[deps.DiffResults]]
deps = ["StaticArraysCore"]
git-tree-sha1 = "782dd5f4561f5d267313f23853baaaa4c52ea621"
uuid = "163ba53b-c6d8-5494-b064-1a9d43ac40c5"
version = "1.1.0"

[[deps.DiffRules]]
deps = ["IrrationalConstants", "LogExpFunctions", "NaNMath", "Random", "SpecialFunctions"]
git-tree-sha1 = "c5b6685d53f933c11404a3ae9822afe30d522494"
uuid = "b552c78f-8df3-52c6-915a-8e097449b14b"
version = "1.12.2"

[[deps.DifferentialEquations]]
deps = ["BoundaryValueDiffEq", "DelayDiffEq", "DiffEqBase", "DiffEqCallbacks", "DiffEqNoiseProcess", "JumpProcesses", "LinearAlgebra", "LinearSolve", "OrdinaryDiffEq", "Random", "RecursiveArrayTools", "Reexport", "SciMLBase", "SteadyStateDiffEq", "StochasticDiffEq", "Sundials"]
git-tree-sha1 = "418dad2cfd7377a474326bc86a23cb645fac6527"
uuid = "0c46a032-eb83-5123-abaf-570d42b7fbaa"
version = "7.6.0"

[[deps.Distances]]
deps = ["LinearAlgebra", "SparseArrays", "Statistics", "StatsAPI"]
git-tree-sha1 = "3258d0659f812acde79e8a74b11f17ac06d0ca04"
uuid = "b4f34e82-e78d-54a5-968a-f98e89d6e8f7"
version = "0.10.7"

[[deps.Distributed]]
deps = ["Random", "Serialization", "Sockets"]
uuid = "8ba89e20-285c-5b6f-9357-94700520ee1b"

[[deps.Distributions]]
deps = ["ChainRulesCore", "DensityInterface", "FillArrays", "LinearAlgebra", "PDMats", "Printf", "QuadGK", "Random", "SparseArrays", "SpecialFunctions", "Statistics", "StatsBase", "StatsFuns", "Test"]
git-tree-sha1 = "74911ad88921455c6afcad1eefa12bd7b1724631"
uuid = "31c24e10-a181-5473-b8eb-7969acd0382f"
version = "0.25.80"

[[deps.DocStringExtensions]]
deps = ["LibGit2"]
git-tree-sha1 = "2fb1e02f2b635d0845df5d7c167fec4dd739b00d"
uuid = "ffbed154-4ef7-542d-bbb7-c09d3a79fcae"
version = "0.9.3"

[[deps.DomainSets]]
deps = ["CompositeTypes", "IntervalSets", "LinearAlgebra", "Random", "StaticArrays", "Statistics"]
git-tree-sha1 = "988e2db482abeb69efc76ae8b6eba2e93805ee70"
uuid = "5b8099bc-c8ec-5219-889f-1d9e522a28bf"
version = "0.5.15"

[[deps.Downloads]]
deps = ["ArgTools", "FileWatching", "LibCURL", "NetworkOptions"]
uuid = "f43a241f-c20a-4ad4-852c-f6b1247861c6"
version = "1.6.0"

[[deps.DualNumbers]]
deps = ["Calculus", "NaNMath", "SpecialFunctions"]
git-tree-sha1 = "5837a837389fccf076445fce071c8ddaea35a566"
uuid = "fa6b7ba4-c1ee-5f82-b5fc-ecf0adba8f74"
version = "0.6.8"

[[deps.DynamicPolynomials]]
deps = ["DataStructures", "Future", "LinearAlgebra", "MultivariatePolynomials", "MutableArithmetics", "Pkg", "Reexport", "Test"]
git-tree-sha1 = "d0fa82f39c2a5cdb3ee385ad52bc05c42cb4b9f0"
uuid = "7c1d4256-1411-5781-91ec-d7bc3513ac07"
version = "0.4.5"

[[deps.EnumX]]
git-tree-sha1 = "bdb1942cd4c45e3c678fd11569d5cccd80976237"
uuid = "4e289a0a-7415-4d19-859d-a7e5c4648b56"
version = "1.0.4"

[[deps.ExponentialUtilities]]
deps = ["Adapt", "ArrayInterfaceCore", "ArrayInterfaceGPUArrays", "GPUArraysCore", "GenericSchur", "LinearAlgebra", "Printf", "SparseArrays", "libblastrampoline_jll"]
git-tree-sha1 = "9837d3f3a904c7a7ab9337759c0093d3abea1d81"
uuid = "d4d017d3-3776-5f7e-afef-a10c40355c18"
version = "1.22.0"

[[deps.ExprTools]]
git-tree-sha1 = "56559bbef6ca5ea0c0818fa5c90320398a6fbf8d"
uuid = "e2ba6199-217a-4e67-a87a-7c52f15ade04"
version = "0.1.8"

[[deps.FNVHash]]
git-tree-sha1 = "d6de2c735a8bffce9bc481942dfa453cc815357e"
uuid = "5207ad80-27db-4d23-8732-fa0bd339ea89"
version = "0.1.0"

[[deps.FastBroadcast]]
deps = ["ArrayInterface", "ArrayInterfaceCore", "LinearAlgebra", "Polyester", "Static", "StrideArraysCore"]
git-tree-sha1 = "4bef892787c972913d4d84e7255400759bb650e5"
uuid = "7034ab61-46d4-4ed7-9d0f-46aef9175898"
version = "0.2.4"

[[deps.FastClosures]]
git-tree-sha1 = "acebe244d53ee1b461970f8910c235b259e772ef"
uuid = "9aa1b823-49e4-5ca5-8b0f-3971ec8bab6a"
version = "0.3.2"

[[deps.FastLapackInterface]]
deps = ["LinearAlgebra"]
git-tree-sha1 = "7fbaf9f73cd4c8561702ea9b16acf3f99d913fe4"
uuid = "29a986be-02c6-4525-aec4-84b980013641"
version = "1.2.8"

[[deps.FilePathsBase]]
deps = ["Compat", "Dates", "Mmap", "Printf", "Test", "UUIDs"]
git-tree-sha1 = "e27c4ebe80e8699540f2d6c805cc12203b614f12"
uuid = "48062228-2e41-5def-b9a4-89aafe57970f"
version = "0.9.20"

[[deps.FileWatching]]
uuid = "7b1f6079-737a-58dc-b8bc-7a2ca5c1b5ee"

[[deps.FillArrays]]
deps = ["LinearAlgebra", "Random", "SparseArrays", "Statistics"]
git-tree-sha1 = "9a0472ec2f5409db243160a8b030f94c380167a3"
uuid = "1a297f60-69ca-5386-bcde-b61e274b549b"
version = "0.13.6"

[[deps.FiniteDiff]]
deps = ["ArrayInterfaceCore", "LinearAlgebra", "Requires", "Setfield", "SparseArrays", "StaticArrays"]
git-tree-sha1 = "04ed1f0029b6b3af88343e439b995141cb0d0b8d"
uuid = "6a86dc24-6348-571c-b903-95158fe2bd41"
version = "2.17.0"

[[deps.FixedPointNumbers]]
deps = ["Statistics"]
git-tree-sha1 = "335bfdceacc84c5cdf16aadc768aa5ddfc5383cc"
uuid = "53c48c17-4a7d-5ca2-90c5-79b7896eea93"
version = "0.8.4"

[[deps.Formatting]]
deps = ["Printf"]
git-tree-sha1 = "8339d61043228fdd3eb658d86c926cb282ae72a8"
uuid = "59287772-0a20-5a39-b81b-1366585eb4c0"
version = "0.4.2"

[[deps.ForwardDiff]]
deps = ["CommonSubexpressions", "DiffResults", "DiffRules", "LinearAlgebra", "LogExpFunctions", "NaNMath", "Preferences", "Printf", "Random", "SpecialFunctions", "StaticArrays"]
git-tree-sha1 = "a69dd6db8a809f78846ff259298678f0d6212180"
uuid = "f6369f11-7733-5829-9624-2563aa707210"
version = "0.10.34"

[[deps.FunctionWrappers]]
git-tree-sha1 = "d62485945ce5ae9c0c48f124a84998d755bae00e"
uuid = "069b7b12-0de2-55c6-9aab-29f3d0a68a2e"
version = "1.1.3"

[[deps.FunctionWrappersWrappers]]
deps = ["FunctionWrappers"]
git-tree-sha1 = "a5e6e7f12607e90d71b09e6ce2c965e41b337968"
uuid = "77dc65aa-8811-40c2-897b-53d922fa7daf"
version = "0.1.1"

[[deps.Future]]
deps = ["Random"]
uuid = "9fa8497b-333b-5362-9e8d-4d0656e87820"

[[deps.GPUArrays]]
deps = ["Adapt", "GPUArraysCore", "LLVM", "LinearAlgebra", "Printf", "Random", "Reexport", "Serialization", "Statistics"]
git-tree-sha1 = "45d7deaf05cbb44116ba785d147c518ab46352d7"
uuid = "0c68f7d7-f131-5f86-a1c3-88cf8149b2d7"
version = "8.5.0"

[[deps.GPUArraysCore]]
deps = ["Adapt"]
git-tree-sha1 = "6872f5ec8fd1a38880f027a26739d42dcda6691f"
uuid = "46192b85-c4d5-4398-a991-12ede77f4527"
version = "0.1.2"

[[deps.GPUCompiler]]
deps = ["ExprTools", "InteractiveUtils", "LLVM", "Libdl", "Logging", "TimerOutputs", "UUIDs"]
git-tree-sha1 = "48832a7cacbe56e591a7bef690c78b9d00bcc692"
uuid = "61eb1bfa-7361-4325-ad38-22787b887f55"
version = "0.17.1"

[[deps.GenericSchur]]
deps = ["LinearAlgebra", "Printf"]
git-tree-sha1 = "fb69b2a645fa69ba5f474af09221b9308b160ce6"
uuid = "c145ed77-6b09-5dd9-b285-bf645a82121e"
version = "0.5.3"

[[deps.Glob]]
git-tree-sha1 = "4df9f7e06108728ebf00a0a11edee4b29a482bb2"
uuid = "c27321d9-0574-5035-807b-f59d2c89b15c"
version = "1.3.0"

[[deps.Graphs]]
deps = ["ArnoldiMethod", "Compat", "DataStructures", "Distributed", "Inflate", "LinearAlgebra", "Random", "SharedArrays", "SimpleTraits", "SparseArrays", "Statistics"]
git-tree-sha1 = "ba2d094a88b6b287bd25cfa86f301e7693ffae2f"
uuid = "86223c79-3864-5bf0-83f7-82e725a168b6"
version = "1.7.4"

[[deps.Groebner]]
deps = ["AbstractAlgebra", "Combinatorics", "Logging", "MultivariatePolynomials", "Primes", "Random"]
git-tree-sha1 = "47f0f03eddecd7ad59c42b1dd46d5f42916aff63"
uuid = "0b43b601-686d-58a3-8a1c-6623616c7cd4"
version = "0.2.11"

[[deps.GroupsCore]]
deps = ["Markdown", "Random"]
git-tree-sha1 = "9e1a5e9f3b81ad6a5c613d181664a0efc6fe6dd7"
uuid = "d5909c97-4eac-4ecc-a3dc-fdd0858a4120"
version = "0.4.0"

[[deps.HostCPUFeatures]]
deps = ["BitTwiddlingConvenienceFunctions", "IfElse", "Libdl", "Static"]
git-tree-sha1 = "f64b890b2efa4de81520d2b0fbdc9aadb65bdf53"
uuid = "3e5b6fbb-0976-4d2c-9146-d79de83f2fb0"
version = "0.1.13"

[[deps.HypergeometricFunctions]]
deps = ["DualNumbers", "LinearAlgebra", "OpenLibm_jll", "SpecialFunctions", "Test"]
git-tree-sha1 = "709d864e3ed6e3545230601f94e11ebc65994641"
uuid = "34004b35-14d8-5ef3-9330-4cdb6864b03a"
version = "0.3.11"

[[deps.IJulia]]
deps = ["Base64", "Conda", "Dates", "InteractiveUtils", "JSON", "Libdl", "Logging", "Markdown", "MbedTLS", "Pkg", "Printf", "REPL", "Random", "SoftGlobalScope", "Test", "UUIDs", "ZMQ"]
git-tree-sha1 = "59e19713542dd9dd02f31d59edbada69530d6a14"
uuid = "7073ff75-c697-5162-941a-fcdaad2a7d2a"
version = "1.24.0"

[[deps.IfElse]]
git-tree-sha1 = "debdd00ffef04665ccbb3e150747a77560e8fad1"
uuid = "615f187c-cbe4-4ef1-ba3b-2fcf58d6d173"
version = "0.1.1"

[[deps.Inflate]]
git-tree-sha1 = "5cd07aab533df5170988219191dfad0519391428"
uuid = "d25df0c9-e2be-5dd7-82c8-3ad0b3e990b9"
version = "0.1.3"

[[deps.InitialValues]]
git-tree-sha1 = "4da0f88e9a39111c2fa3add390ab15f3a44f3ca3"
uuid = "22cec73e-a1b8-11e9-2c92-598750a2cf9c"
version = "0.3.1"

[[deps.InlineStrings]]
deps = ["Parsers"]
git-tree-sha1 = "0cf92ec945125946352f3d46c96976ab972bde6f"
uuid = "842dd82b-1e85-43dc-bf29-5d0ee9dffc48"
version = "1.3.2"

[[deps.IntegerMathUtils]]
git-tree-sha1 = "f366daebdfb079fd1fe4e3d560f99a0c892e15bc"
uuid = "18e54dd8-cb9d-406c-a71d-865a43cbb235"
version = "0.1.0"

[[deps.InteractiveUtils]]
deps = ["Markdown"]
uuid = "b77e0a4c-d291-57a0-90e8-8db25a27a240"

[[deps.Interpolations]]
deps = ["Adapt", "AxisAlgorithms", "ChainRulesCore", "LinearAlgebra", "OffsetArrays", "Random", "Ratios", "Requires", "SharedArrays", "SparseArrays", "StaticArrays", "WoodburyMatrices"]
git-tree-sha1 = "721ec2cf720536ad005cb38f50dbba7b02419a15"
uuid = "a98d9a8b-a2ab-59e6-89dd-64a1c18fca59"
version = "0.14.7"

[[deps.IntervalSets]]
deps = ["Dates", "Random", "Statistics"]
git-tree-sha1 = "3f91cd3f56ea48d4d2a75c2a65455c5fc74fa347"
uuid = "8197267c-284f-5f27-9208-e0e47529a953"
version = "0.7.3"

[[deps.InverseFunctions]]
deps = ["Test"]
git-tree-sha1 = "49510dfcb407e572524ba94aeae2fced1f3feb0f"
uuid = "3587e190-3f89-42d0-90ee-14403ec27112"
version = "0.1.8"

[[deps.InvertedIndices]]
git-tree-sha1 = "82aec7a3dd64f4d9584659dc0b62ef7db2ef3e19"
uuid = "41ab1584-1d38-5bbf-9106-f11c6c58b48f"
version = "1.2.0"

[[deps.IrrationalConstants]]
git-tree-sha1 = "7fd44fd4ff43fc60815f8e764c0f352b83c49151"
uuid = "92d709cd-6900-40b7-9082-c6be49f344b6"
version = "0.1.1"

[[deps.IterativeSolvers]]
deps = ["LinearAlgebra", "Printf", "Random", "RecipesBase", "SparseArrays"]
git-tree-sha1 = "1169632f425f79429f245113b775a0e3d121457c"
uuid = "42fd0dbc-a981-5370-80f2-aaf504508153"
version = "0.9.2"

[[deps.IteratorInterfaceExtensions]]
git-tree-sha1 = "a3f24677c21f5bbe9d2a714f95dcd58337fb2856"
uuid = "82899510-4779-5014-852e-03e436cf321d"
version = "1.0.0"

[[deps.JLLWrappers]]
deps = ["Preferences"]
git-tree-sha1 = "abc9885a7ca2052a736a600f7fa66209f96506e1"
uuid = "692b3bcd-3c85-4b1f-b108-f13ce0eb3210"
version = "1.4.1"

[[deps.JSON]]
deps = ["Dates", "Mmap", "Parsers", "Unicode"]
git-tree-sha1 = "3c837543ddb02250ef42f4738347454f95079d4e"
uuid = "682c06a0-de6a-54ab-a142-c8b1cf79cde6"
version = "0.21.3"

[[deps.JSON3]]
deps = ["Dates", "Mmap", "Parsers", "SnoopPrecompile", "StructTypes", "UUIDs"]
git-tree-sha1 = "84b10656a41ef564c39d2d477d7236966d2b5683"
uuid = "0f8b85d8-7281-11e9-16c2-39a750bddbf1"
version = "1.12.0"

[[deps.JuliaFormatter]]
deps = ["CSTParser", "CommonMark", "DataStructures", "Glob", "Pkg", "Tokenize"]
git-tree-sha1 = "447f5702b5271d99f064e924061bc05e26f52c4c"
uuid = "98e50ef6-434e-11e9-1051-2b60c6c9e899"
version = "1.0.18"

[[deps.JumpProcesses]]
deps = ["ArrayInterfaceCore", "DataStructures", "DiffEqBase", "DocStringExtensions", "FunctionWrappers", "Graphs", "LinearAlgebra", "Markdown", "PoissonRandom", "Random", "RandomNumbers", "RecursiveArrayTools", "Reexport", "SciMLBase", "StaticArrays", "TreeViews", "UnPack"]
git-tree-sha1 = "09ed2720b2e343e48780a3156c4a6cef8dd54192"
uuid = "ccbc3e58-028d-4f4c-8cd5-9ae44345cda5"
version = "9.3.1"

[[deps.KLU]]
deps = ["LinearAlgebra", "SparseArrays", "SuiteSparse_jll"]
git-tree-sha1 = "764164ed65c30738750965d55652db9c94c59bfe"
uuid = "ef3ab10e-7fda-4108-b977-705223b18434"
version = "0.4.0"

[[deps.KernelAbstractions]]
deps = ["Adapt", "Atomix", "InteractiveUtils", "LinearAlgebra", "MacroTools", "SparseArrays", "StaticArrays", "UUIDs", "UnsafeAtomics", "UnsafeAtomicsLLVM"]
git-tree-sha1 = "cf9cae1c4c1ff83f6c02cfaf01698f05448e8325"
uuid = "63c18a36-062a-441e-b654-da1e3ab1ce7c"
version = "0.8.6"

[[deps.Krylov]]
deps = ["LinearAlgebra", "Printf", "SparseArrays"]
git-tree-sha1 = "dd90aacbfb622f898a97c2a4411ac49101ebab8a"
uuid = "ba0b0d4f-ebba-5204-a429-3ac8c609bfb7"
version = "0.9.0"

[[deps.KrylovKit]]
deps = ["ChainRulesCore", "GPUArraysCore", "LinearAlgebra", "Printf"]
git-tree-sha1 = "1a5e1d9941c783b0119897d29f2eb665d876ecf3"
uuid = "0b1a1467-8014-51b9-945f-bf0ae24f4b77"
version = "0.6.0"

[[deps.LLVM]]
deps = ["CEnum", "LLVMExtra_jll", "Libdl", "Printf", "Unicode"]
git-tree-sha1 = "088dd02b2797f0233d92583562ab669de8517fd1"
uuid = "929cbde3-209d-540e-8aea-75f648917ca0"
version = "4.14.1"

[[deps.LLVMExtra_jll]]
deps = ["Artifacts", "JLLWrappers", "LazyArtifacts", "Libdl", "Pkg", "TOML"]
git-tree-sha1 = "7718cf44439c676bc0ec66a87099f41015a522d6"
uuid = "dad2f222-ce93-54a1-a47d-0025e8a3acab"
version = "0.0.16+2"

[[deps.LZO_jll]]
deps = ["Artifacts", "JLLWrappers", "Libdl", "Pkg"]
git-tree-sha1 = "e5b909bcf985c5e2605737d2ce278ed791b89be6"
uuid = "dd4b983a-f0e5-5f8d-a1b7-129d4a5fb1ac"
version = "2.10.1+0"

[[deps.LaTeXStrings]]
git-tree-sha1 = "f2355693d6778a178ade15952b7ac47a4ff97996"
uuid = "b964fa9f-0449-5b57-a5c2-d3ea65f4040f"
version = "1.3.0"

[[deps.LabelledArrays]]
deps = ["ArrayInterfaceCore", "ArrayInterfaceStaticArrays", "ArrayInterfaceStaticArraysCore", "ChainRulesCore", "ForwardDiff", "LinearAlgebra", "MacroTools", "PreallocationTools", "RecursiveArrayTools", "StaticArrays"]
git-tree-sha1 = "dae002226b59701dbafd7e2dd757df1bd83442fd"
uuid = "2ee39098-c373-598a-b85f-a56591580800"
version = "1.12.5"

[[deps.LambertW]]
git-tree-sha1 = "2d9f4009c486ef676646bca06419ac02061c088e"
uuid = "984bce1d-4616-540c-a9ee-88d1112d94c9"
version = "0.4.5"

[[deps.Latexify]]
deps = ["Formatting", "InteractiveUtils", "LaTeXStrings", "MacroTools", "Markdown", "OrderedCollections", "Printf", "Requires"]
git-tree-sha1 = "2422f47b34d4b127720a18f86fa7b1aa2e141f29"
uuid = "23fbe1c1-3f47-55db-b15f-69d7ec21a316"
version = "0.15.18"

[[deps.LayoutPointers]]
deps = ["ArrayInterface", "ArrayInterfaceOffsetArrays", "ArrayInterfaceStaticArrays", "LinearAlgebra", "ManualMemory", "SIMDTypes", "Static"]
git-tree-sha1 = "7e34177793212f6d64d045ee47d2883f09fffacc"
uuid = "10f19ff3-798f-405d-979b-55457f8fc047"
version = "0.1.12"

[[deps.LazyArrays]]
deps = ["ArrayLayouts", "FillArrays", "LinearAlgebra", "MacroTools", "MatrixFactorizations", "SparseArrays", "StaticArrays"]
git-tree-sha1 = "f6742270b551e03bd8ba64e1c754f5fab089ab63"
uuid = "5078a376-72f3-5289-bfd5-ec5146d43c02"
version = "0.22.16"

[[deps.LazyArtifacts]]
deps = ["Artifacts", "Pkg"]
uuid = "4af54fe1-eca0-43a8-85a7-787d91b784e3"

[[deps.LevyArea]]
deps = ["LinearAlgebra", "Random", "SpecialFunctions"]
git-tree-sha1 = "56513a09b8e0ae6485f34401ea9e2f31357958ec"
uuid = "2d8b4e74-eb68-11e8-0fb9-d5eb67b50637"
version = "1.0.0"

[[deps.LibCURL]]
deps = ["LibCURL_jll", "MozillaCACerts_jll"]
uuid = "b27032c2-a3e7-50c8-80cd-2d36dbcbfd21"
version = "0.6.3"

[[deps.LibCURL_jll]]
deps = ["Artifacts", "LibSSH2_jll", "Libdl", "MbedTLS_jll", "Zlib_jll", "nghttp2_jll"]
uuid = "deac9b47-8bc7-5906-a0fe-35ac56dc84c0"
version = "7.84.0+0"

[[deps.LibGit2]]
deps = ["Base64", "NetworkOptions", "Printf", "SHA"]
uuid = "76f85450-5226-5b5a-8eaa-529ad045b433"

[[deps.LibSSH2_jll]]
deps = ["Artifacts", "Libdl", "MbedTLS_jll"]
uuid = "29816b5a-b9ab-546f-933c-edad1886dfa8"
version = "1.10.2+0"

[[deps.Libdl]]
uuid = "8f399da3-3557-5675-b5ff-fb832c97cbdb"

[[deps.LightBSON]]
deps = ["DataStructures", "Dates", "DecFP", "FNVHash", "JSON3", "Sockets", "StructTypes", "Transducers", "UUIDs", "UnsafeArrays", "WeakRefStrings"]
git-tree-sha1 = "66369db4570bcd852bde2dd39beaa559bc9890dd"
uuid = "a4a7f996-b3a6-4de6-b9db-2fa5f350df41"
version = "0.2.16"

[[deps.LineSearches]]
deps = ["LinearAlgebra", "NLSolversBase", "NaNMath", "Parameters", "Printf"]
git-tree-sha1 = "7bbea35cec17305fc70a0e5b4641477dc0789d9d"
uuid = "d3d80556-e9d4-5f37-9878-2ab0fcc64255"
version = "7.2.0"

[[deps.LinearAlgebra]]
deps = ["Libdl", "libblastrampoline_jll"]
uuid = "37e2e46d-f89d-539d-b4ee-838fcccc9c8e"

[[deps.LinearSolve]]
deps = ["ArrayInterfaceCore", "DocStringExtensions", "FastLapackInterface", "GPUArraysCore", "IterativeSolvers", "KLU", "Krylov", "KrylovKit", "LinearAlgebra", "Preferences", "RecursiveFactorization", "Reexport", "SciMLBase", "Setfield", "SnoopPrecompile", "SparseArrays", "Sparspak", "SuiteSparse", "UnPack"]
git-tree-sha1 = "cf1227e369513687658476e466a5b73a7c3dfa1f"
uuid = "7ed4a6bd-45f5-4d41-b270-4a48e9bafcae"
version = "1.33.0"

[[deps.LogExpFunctions]]
deps = ["ChainRulesCore", "ChangesOfVariables", "DocStringExtensions", "InverseFunctions", "IrrationalConstants", "LinearAlgebra"]
git-tree-sha1 = "946607f84feb96220f480e0422d3484c49c00239"
uuid = "2ab3a3ac-af41-5b50-aa03-7779005ae688"
version = "0.3.19"

[[deps.Logging]]
uuid = "56ddb016-857b-54e1-b83d-db4d58db5568"

[[deps.LoggingExtras]]
deps = ["Dates", "Logging"]
git-tree-sha1 = "cedb76b37bc5a6c702ade66be44f831fa23c681e"
uuid = "e6f89c97-d47a-5376-807f-9c37f3926c36"
version = "1.0.0"

[[deps.LoopVectorization]]
deps = ["ArrayInterface", "ArrayInterfaceCore", "ArrayInterfaceOffsetArrays", "ArrayInterfaceStaticArrays", "CPUSummary", "ChainRulesCore", "CloseOpenIntervals", "DocStringExtensions", "ForwardDiff", "HostCPUFeatures", "IfElse", "LayoutPointers", "LinearAlgebra", "OffsetArrays", "PolyesterWeave", "SIMDDualNumbers", "SIMDTypes", "SLEEFPirates", "SnoopPrecompile", "SpecialFunctions", "Static", "ThreadingUtilities", "UnPack", "VectorizationBase"]
git-tree-sha1 = "155132d68bc33c826dbdeb452c5d0a79e2d0e586"
uuid = "bdcacae8-1622-11e9-2a5c-532679323890"
version = "0.12.146"

[[deps.Lz4_jll]]
deps = ["Artifacts", "JLLWrappers", "Libdl", "Pkg"]
git-tree-sha1 = "5d494bc6e85c4c9b626ee0cab05daa4085486ab1"
uuid = "5ced341a-0733-55b8-9ab6-a4889d929147"
version = "1.9.3+0"

[[deps.MacroTools]]
deps = ["Markdown", "Random"]
git-tree-sha1 = "42324d08725e200c23d4dfb549e0d5d89dede2d2"
uuid = "1914dd2f-81c6-5fcd-8719-6d5c9610ff09"
version = "0.5.10"

[[deps.ManualMemory]]
git-tree-sha1 = "bcaef4fc7a0cfe2cba636d84cda54b5e4e4ca3cd"
uuid = "d125e4d3-2237-4719-b19c-fa641b8a4667"
version = "0.1.8"

[[deps.Markdown]]
deps = ["Base64"]
uuid = "d6f4376e-aef5-505a-96c1-9c027394607a"

[[deps.MatrixFactorizations]]
deps = ["ArrayLayouts", "LinearAlgebra", "Printf", "Random"]
git-tree-sha1 = "22cf8c4315c417300b753c0def0fcb94e83d79b1"
uuid = "a3b82374-2e81-5b9e-98ce-41277c0e4c87"
version = "0.9.5"

[[deps.MbedTLS]]
deps = ["Dates", "MbedTLS_jll", "MozillaCACerts_jll", "Random", "Sockets"]
git-tree-sha1 = "03a9b9718f5682ecb107ac9f7308991db4ce395b"
uuid = "739be429-bea8-5141-9913-cc70e7f3736d"
version = "1.1.7"

[[deps.MbedTLS_jll]]
deps = ["Artifacts", "Libdl"]
uuid = "c8ffd9c3-330d-5841-b78e-0817d7145fa1"
version = "2.28.0+0"

[[deps.Metatheory]]
deps = ["AutoHashEquals", "DataStructures", "Dates", "DocStringExtensions", "Parameters", "Reexport", "TermInterface", "ThreadsX", "TimerOutputs"]
git-tree-sha1 = "0f39bc7f71abdff12ead4fc4a7d998fb2f3c171f"
uuid = "e9d8d322-4543-424a-9be4-0cc815abe26c"
version = "1.3.5"

[[deps.MethodOfLines]]
deps = ["Combinatorics", "DiffEqBase", "DomainSets", "IfElse", "Interpolations", "Latexify", "LinearAlgebra", "ModelingToolkit", "OrdinaryDiffEq", "SciMLBase", "StaticArrays", "SymbolicUtils", "Symbolics", "TermInterface"]
git-tree-sha1 = "863017fd2d2e30e9505bb1b08ba722cbf48bb58a"
uuid = "94925ecb-adb7-4558-8ed8-f975c56a0bf4"
version = "0.8.1"

[[deps.MicroCollections]]
deps = ["BangBang", "InitialValues", "Setfield"]
git-tree-sha1 = "4d5917a26ca33c66c8e5ca3247bd163624d35493"
uuid = "128add7d-3638-4c79-886c-908ea0c25c34"
version = "0.1.3"

[[deps.Missings]]
deps = ["DataAPI"]
git-tree-sha1 = "f66bdc5de519e8f8ae43bdc598782d35a25b1272"
uuid = "e1d29d7a-bbdc-5cf2-9ac0-f12de2c33e28"
version = "1.1.0"

[[deps.Mmap]]
uuid = "a63ad114-7e13-5084-954f-fe012c677804"

[[deps.Mocking]]
deps = ["Compat", "ExprTools"]
git-tree-sha1 = "c272302b22479a24d1cf48c114ad702933414f80"
uuid = "78c3b35d-d492-501b-9361-3d52fe80e533"
version = "0.7.5"

[[deps.ModelingToolkit]]
deps = ["AbstractTrees", "ArrayInterfaceCore", "Combinatorics", "Compat", "ConstructionBase", "DataStructures", "DiffEqBase", "DiffEqCallbacks", "DiffRules", "Distributed", "Distributions", "DocStringExtensions", "DomainSets", "ForwardDiff", "FunctionWrappersWrappers", "Graphs", "IfElse", "InteractiveUtils", "JuliaFormatter", "JumpProcesses", "LabelledArrays", "Latexify", "Libdl", "LinearAlgebra", "MacroTools", "NaNMath", "RecursiveArrayTools", "Reexport", "RuntimeGeneratedFunctions", "SciMLBase", "Serialization", "Setfield", "SimpleNonlinearSolve", "SparseArrays", "SpecialFunctions", "StaticArrays", "SymbolicIndexingInterface", "SymbolicUtils", "Symbolics", "UnPack", "Unitful"]
git-tree-sha1 = "86b5a6f7c95483eb8d98f7861e3f7c90113202b5"
uuid = "961ee093-0014-501f-94e3-6117800e7a78"
version = "8.40.0"

[[deps.MozillaCACerts_jll]]
uuid = "14a3606d-f60d-562e-9121-12d972cd8159"
version = "2022.2.1"

[[deps.MuladdMacro]]
git-tree-sha1 = "cac9cc5499c25554cba55cd3c30543cff5ca4fab"
uuid = "46d2c3a1-f734-5fdb-9937-b9b9aeba4221"
version = "0.2.4"

[[deps.MultivariatePolynomials]]
deps = ["ChainRulesCore", "DataStructures", "LinearAlgebra", "MutableArithmetics"]
git-tree-sha1 = "393fc4d82a73c6fe0e2963dd7c882b09257be537"
uuid = "102ac46a-7ee4-5c85-9060-abc95bfdeaa3"
version = "0.4.6"

[[deps.MutableArithmetics]]
deps = ["LinearAlgebra", "SparseArrays", "Test"]
git-tree-sha1 = "aa532179d4a643d4bd9f328589ca01fa20a0d197"
uuid = "d8a4904e-b15c-11e9-3269-09a3773c0cb0"
version = "1.1.0"

[[deps.NLSolversBase]]
deps = ["DiffResults", "Distributed", "FiniteDiff", "ForwardDiff"]
git-tree-sha1 = "a0b464d183da839699f4c79e7606d9d186ec172c"
uuid = "d41bc354-129a-5804-8e4c-c37616107c6c"
version = "7.8.3"

[[deps.NLsolve]]
deps = ["Distances", "LineSearches", "LinearAlgebra", "NLSolversBase", "Printf", "Reexport"]
git-tree-sha1 = "019f12e9a1a7880459d0173c182e6a99365d7ac1"
uuid = "2774e3e8-f4cf-5e23-947b-6d7e65073b56"
version = "4.5.1"

[[deps.NaNMath]]
deps = ["OpenLibm_jll"]
git-tree-sha1 = "a7c3d1da1189a1c2fe843a3bfa04d18d20eb3211"
uuid = "77ba4419-2d1f-58cd-9bb1-8ffee604a2e3"
version = "1.0.1"

[[deps.NetworkOptions]]
uuid = "ca575930-c2e3-43a9-ace4-1e988b2c1908"
version = "1.2.0"

[[deps.NonlinearSolve]]
deps = ["ArrayInterfaceCore", "DiffEqBase", "FiniteDiff", "ForwardDiff", "LinearAlgebra", "LinearSolve", "RecursiveArrayTools", "Reexport", "SciMLBase", "SimpleNonlinearSolve", "SnoopPrecompile", "SparseArrays", "SparseDiffTools", "StaticArraysCore", "UnPack"]
git-tree-sha1 = "7142ca5ab9bd7452cafb29f7d51f574a09d69052"
uuid = "8913a72c-1f9b-4ce2-8d82-65094dcecaec"
version = "1.1.1"

[[deps.OffsetArrays]]
deps = ["Adapt"]
git-tree-sha1 = "f71d8950b724e9ff6110fc948dff5a329f901d64"
uuid = "6fe1bfb0-de20-5000-8ca7-80f57d26f881"
version = "1.12.8"

[[deps.OpenBLAS_jll]]
deps = ["Artifacts", "CompilerSupportLibraries_jll", "Libdl"]
uuid = "4536629a-c528-5b80-bd46-f80d51c5b363"
version = "0.3.20+0"

[[deps.OpenLibm_jll]]
deps = ["Artifacts", "Libdl"]
uuid = "05823500-19ac-5b8b-9628-191a04bc5112"
version = "0.8.1+0"

[[deps.OpenSpecFun_jll]]
deps = ["Artifacts", "CompilerSupportLibraries_jll", "JLLWrappers", "Libdl", "Pkg"]
git-tree-sha1 = "13652491f6856acfd2db29360e1bbcd4565d04f1"
uuid = "efe28fd5-8261-553b-a9e1-b2916fc3738e"
version = "0.5.5+0"

[[deps.Optim]]
deps = ["Compat", "FillArrays", "ForwardDiff", "LineSearches", "LinearAlgebra", "NLSolversBase", "NaNMath", "Parameters", "PositiveFactorizations", "Printf", "SparseArrays", "StatsBase"]
git-tree-sha1 = "1903afc76b7d01719d9c30d3c7d501b61db96721"
uuid = "429524aa-4258-5aef-a3af-852621145aeb"
version = "1.7.4"

[[deps.OrderedCollections]]
git-tree-sha1 = "85f8e6578bf1f9ee0d11e7bb1b1456435479d47c"
uuid = "bac558e1-5e72-5ebc-8fee-abe8a469f55d"
version = "1.4.1"

[[deps.OrdinaryDiffEq]]
deps = ["Adapt", "ArrayInterface", "ArrayInterfaceCore", "ArrayInterfaceGPUArrays", "ArrayInterfaceStaticArrays", "ArrayInterfaceStaticArraysCore", "DataStructures", "DiffEqBase", "DocStringExtensions", "ExponentialUtilities", "FastBroadcast", "FastClosures", "FiniteDiff", "ForwardDiff", "FunctionWrappersWrappers", "LinearAlgebra", "LinearSolve", "Logging", "LoopVectorization", "MacroTools", "MuladdMacro", "NLsolve", "NonlinearSolve", "Polyester", "PreallocationTools", "Preferences", "RecursiveArrayTools", "Reexport", "SciMLBase", "SciMLNLSolve", "SimpleNonlinearSolve", "SnoopPrecompile", "SparseArrays", "SparseDiffTools", "StaticArrays", "UnPack"]
git-tree-sha1 = "ca0c8939dbd3617ae3fdca13374d0b7501a2dd28"
uuid = "1dea7af3-3e70-54e6-95c3-0bf5283fa5ed"
version = "6.37.0"

[[deps.PDMats]]
deps = ["LinearAlgebra", "SparseArrays", "SuiteSparse"]
git-tree-sha1 = "cf494dca75a69712a72b80bc48f59dcf3dea63ec"
uuid = "90014a1f-27ba-587c-ab20-58faa44d9150"
version = "0.11.16"

[[deps.Parameters]]
deps = ["OrderedCollections", "UnPack"]
git-tree-sha1 = "34c0e9ad262e5f7fc75b10a9952ca7692cfc5fbe"
uuid = "d96e819e-fc66-5662-9728-84c9c7592b0a"
version = "0.12.3"

[[deps.Parquet2]]
deps = ["AbstractTrees", "BitIntegers", "CodecLz4", "CodecZlib", "CodecZstd", "DataAPI", "Dates", "DecFP", "FilePathsBase", "FillArrays", "JSON3", "LazyArrays", "LightBSON", "Mmap", "OrderedCollections", "PooledArrays", "SentinelArrays", "Snappy", "StaticArrays", "TableOperations", "Tables", "Thrift", "Transducers", "UUIDs", "WeakRefStrings"]
git-tree-sha1 = "6ccd331e8cee949a96a3b797b0784ad9363c22f5"
uuid = "98572fba-bba0-415d-956f-fa77e587d26d"
version = "0.2.7"

[[deps.Parsers]]
deps = ["Dates", "SnoopPrecompile"]
git-tree-sha1 = "8175fc2b118a3755113c8e68084dc1a9e63c61ee"
uuid = "69de0a69-1ddd-5017-9359-2bf0b02dc9f0"
version = "2.5.3"

[[deps.Pkg]]
deps = ["Artifacts", "Dates", "Downloads", "LibGit2", "Libdl", "Logging", "Markdown", "Printf", "REPL", "Random", "SHA", "Serialization", "TOML", "Tar", "UUIDs", "p7zip_jll"]
uuid = "44cfe95a-1eb2-52ea-b672-e2afdf69b78f"
version = "1.8.0"

[[deps.PoissonRandom]]
deps = ["Random"]
git-tree-sha1 = "45f9da1ceee5078267eb273d065e8aa2f2515790"
uuid = "e409e4f3-bfea-5376-8464-e040bb5c01ab"
version = "0.4.3"

[[deps.Polyester]]
deps = ["ArrayInterface", "BitTwiddlingConvenienceFunctions", "CPUSummary", "IfElse", "ManualMemory", "PolyesterWeave", "Requires", "Static", "StrideArraysCore", "ThreadingUtilities"]
git-tree-sha1 = "7f8dd47630b265df9e1d117137ee1894b195e032"
uuid = "f517fe37-dbe3-4b94-8317-1923a5111588"
version = "0.7.1"

[[deps.PolyesterWeave]]
deps = ["BitTwiddlingConvenienceFunctions", "CPUSummary", "IfElse", "Static", "ThreadingUtilities"]
git-tree-sha1 = "5d0a598c95f67ee0787723e38745cb954d143684"
uuid = "1d0040c9-8b98-4ee7-8388-3f51789ca0ad"
version = "0.2.0"

[[deps.PooledArrays]]
deps = ["DataAPI", "Future"]
git-tree-sha1 = "a6062fe4063cdafe78f4a0a81cfffb89721b30e7"
uuid = "2dfb63ee-cc39-5dd5-95bd-886bf059d720"
version = "1.4.2"

[[deps.PositiveFactorizations]]
deps = ["LinearAlgebra"]
git-tree-sha1 = "17275485f373e6673f7e7f97051f703ed5b15b20"
uuid = "85a6dd25-e78a-55b7-8502-1745935b8125"
version = "0.2.4"

[[deps.PreallocationTools]]
deps = ["Adapt", "ArrayInterfaceCore", "ForwardDiff"]
git-tree-sha1 = "758f3283aba57c53960c8e1900b4c724bf24ba74"
uuid = "d236fae5-4411-538c-8e31-a6e3d9e00b46"
version = "0.4.8"

[[deps.PrecompileTools]]
deps = ["Preferences"]
git-tree-sha1 = "03b4c25b43cb84cee5c90aa9b5ea0a78fd848d2f"
uuid = "aea7be01-6a6a-4083-8856-8a6e6704d82a"
version = "1.2.0"

[[deps.Preferences]]
deps = ["TOML"]
git-tree-sha1 = "47e5f437cc0e7ef2ce8406ce1e7e24d44915f88d"
uuid = "21216c6a-2e73-6563-6e65-726566657250"
version = "1.3.0"

[[deps.PrettyTables]]
deps = ["Crayons", "Formatting", "LaTeXStrings", "Markdown", "Reexport", "StringManipulation", "Tables"]
git-tree-sha1 = "96f6db03ab535bdb901300f88335257b0018689d"
uuid = "08abe8d2-0d0c-5749-adfa-8a2ac140af0d"
version = "2.2.2"

[[deps.Primes]]
deps = ["IntegerMathUtils"]
git-tree-sha1 = "311a2aa90a64076ea0fac2ad7492e914e6feeb81"
uuid = "27ebfcd6-29c5-5fa9-bf4b-fb8fc14df3ae"
version = "0.5.3"

[[deps.Printf]]
deps = ["Unicode"]
uuid = "de0858da-6303-5e67-8744-51eddeeeb8d7"

[[deps.Profile]]
deps = ["Printf"]
uuid = "9abbd945-dff8-562f-b5e8-e1ebf5ef1b79"

[[deps.QuadGK]]
deps = ["DataStructures", "LinearAlgebra"]
git-tree-sha1 = "97aa253e65b784fd13e83774cadc95b38011d734"
uuid = "1fd47b50-473d-5c70-9696-f719f8f3bcdc"
version = "2.6.0"

[[deps.REPL]]
deps = ["InteractiveUtils", "Markdown", "Sockets", "Unicode"]
uuid = "3fa0cd96-eef1-5676-8a61-b3b8758bbffb"

[[deps.Random]]
deps = ["SHA", "Serialization"]
uuid = "9a3f8284-a2c9-5f02-9a11-845980a1fd5c"

[[deps.Random123]]
deps = ["Random", "RandomNumbers"]
git-tree-sha1 = "7a1a306b72cfa60634f03a911405f4e64d1b718b"
uuid = "74087812-796a-5b5d-8853-05524746bad3"
version = "1.6.0"

[[deps.RandomExtensions]]
deps = ["Random", "SparseArrays"]
git-tree-sha1 = "062986376ce6d394b23d5d90f01d81426113a3c9"
uuid = "fb686558-2515-59ef-acaa-46db3789a887"
version = "0.4.3"

[[deps.RandomNumbers]]
deps = ["Random", "Requires"]
git-tree-sha1 = "043da614cc7e95c703498a491e2c21f58a2b8111"
uuid = "e6cf234a-135c-5ec9-84dd-332b85af5143"
version = "1.5.3"

[[deps.Ratios]]
deps = ["Requires"]
git-tree-sha1 = "dc84268fe0e3335a62e315a3a7cf2afa7178a734"
uuid = "c84ed2f1-dad5-54f0-aa8e-dbefe2724439"
version = "0.4.3"

[[deps.RecipesBase]]
deps = ["SnoopPrecompile"]
git-tree-sha1 = "261dddd3b862bd2c940cf6ca4d1c8fe593e457c8"
uuid = "3cdcf5f2-1ef4-517c-9805-6587b60abb01"
version = "1.3.3"

[[deps.RecursiveArrayTools]]
deps = ["Adapt", "ArrayInterfaceCore", "ArrayInterfaceStaticArraysCore", "ChainRulesCore", "DocStringExtensions", "FillArrays", "GPUArraysCore", "IteratorInterfaceExtensions", "LinearAlgebra", "RecipesBase", "StaticArraysCore", "Statistics", "SymbolicIndexingInterface", "Tables", "ZygoteRules"]
git-tree-sha1 = "fcf0962b399f3bc0fa13ae7274db7879c3ef9f1e"
uuid = "731186ca-8d62-57ce-b412-fbd966d074cd"
version = "2.35.0"

[[deps.RecursiveFactorization]]
deps = ["LinearAlgebra", "LoopVectorization", "Polyester", "SnoopPrecompile", "StrideArraysCore", "TriangularSolve"]
git-tree-sha1 = "0ab70c6cd81764d0047921cfc42741d2c2ff402e"
uuid = "f2c3362d-daeb-58d1-803e-2bc74f2840b4"
version = "0.2.15"

[[deps.Reexport]]
git-tree-sha1 = "45e428421666073eab6f2da5c9d310d99bb12f9b"
uuid = "189a3867-3050-52da-a836-e630ba90ab69"
version = "1.2.2"

[[deps.Referenceables]]
deps = ["Adapt"]
git-tree-sha1 = "e681d3bfa49cd46c3c161505caddf20f0e62aaa9"
uuid = "42d2dcc6-99eb-4e98-b66c-637b7d73030e"
version = "0.1.2"

[[deps.Requires]]
deps = ["UUIDs"]
git-tree-sha1 = "838a3a4188e2ded87a4f9f184b4b0d78a1e91cb7"
uuid = "ae029012-a4dd-5104-9daa-d747884805df"
version = "1.3.0"

[[deps.ResettableStacks]]
deps = ["StaticArrays"]
git-tree-sha1 = "256eeeec186fa7f26f2801732774ccf277f05db9"
uuid = "ae5879a3-cd67-5da8-be7f-38c6eb64a37b"
version = "1.1.1"

[[deps.Rmath]]
deps = ["Random", "Rmath_jll"]
git-tree-sha1 = "bf3188feca147ce108c76ad82c2792c57abe7b1f"
uuid = "79098fc4-a85e-5d69-aa6a-4863f24498fa"
version = "0.7.0"

[[deps.Rmath_jll]]
deps = ["Artifacts", "JLLWrappers", "Libdl", "Pkg"]
git-tree-sha1 = "68db32dff12bb6127bac73c209881191bf0efbb7"
uuid = "f50d1b31-88e8-58de-be2c-1cc44531875f"
version = "0.3.0+0"

[[deps.RuntimeGeneratedFunctions]]
deps = ["ExprTools", "SHA", "Serialization"]
git-tree-sha1 = "50314d2ef65fce648975a8e80ae6d8409ebbf835"
uuid = "7e49a35a-f44a-4d26-94aa-eba1b4ca6b47"
version = "0.5.5"

[[deps.SHA]]
uuid = "ea8e919c-243c-51af-8825-aaa63cd721ce"
version = "0.7.0"

[[deps.SIMDDualNumbers]]
deps = ["ForwardDiff", "IfElse", "SLEEFPirates", "VectorizationBase"]
git-tree-sha1 = "dd4195d308df24f33fb10dde7c22103ba88887fa"
uuid = "3cdde19b-5bb0-4aaf-8931-af3e248e098b"
version = "0.1.1"

[[deps.SIMDTypes]]
git-tree-sha1 = "330289636fb8107c5f32088d2741e9fd7a061a5c"
uuid = "94e857df-77ce-4151-89e5-788b33177be4"
version = "0.1.0"

[[deps.SLEEFPirates]]
deps = ["IfElse", "Static", "VectorizationBase"]
git-tree-sha1 = "c8679919df2d3c71f74451321f1efea6433536cc"
uuid = "476501e8-09a2-5ece-8869-fb82de89a1fa"
version = "0.6.37"

[[deps.SciMLBase]]
deps = ["ArrayInterfaceCore", "CommonSolve", "ConstructionBase", "Distributed", "DocStringExtensions", "EnumX", "FunctionWrappersWrappers", "IteratorInterfaceExtensions", "LinearAlgebra", "Logging", "Markdown", "Preferences", "RecipesBase", "RecursiveArrayTools", "RuntimeGeneratedFunctions", "StaticArraysCore", "Statistics", "Tables"]
git-tree-sha1 = "fe89a8113ea445bcff9ee570077830674babb534"
uuid = "0bca4576-84f4-4d90-8ffe-ffa030f20462"
version = "1.81.0"

[[deps.SciMLNLSolve]]
deps = ["LineSearches", "NLsolve", "Reexport", "SciMLBase"]
git-tree-sha1 = "b35d1f5d8afeee44e24915bb767e34fae867502f"
uuid = "e9a6253c-8580-4d32-9898-8661bb511710"
version = "0.1.1"

[[deps.Scratch]]
deps = ["Dates"]
git-tree-sha1 = "f94f779c94e58bf9ea243e77a37e16d9de9126bd"
uuid = "6c6a2e73-6563-6170-7368-637461726353"
version = "1.1.1"

[[deps.SentinelArrays]]
deps = ["Dates", "Random"]
git-tree-sha1 = "efd23b378ea5f2db53a55ae53d3133de4e080aa9"
uuid = "91c51154-3ec4-41a3-a24f-3f23e20d615c"
version = "1.3.16"

[[deps.Serialization]]
uuid = "9e88b42a-f829-5b0c-bbe9-9e923198166b"

[[deps.Setfield]]
deps = ["ConstructionBase", "Future", "MacroTools", "StaticArraysCore"]
git-tree-sha1 = "e2cc6d8c88613c05e1defb55170bf5ff211fbeac"
uuid = "efcf1570-3423-57d1-acb7-fd33fddbac46"
version = "1.1.1"

[[deps.SharedArrays]]
deps = ["Distributed", "Mmap", "Random", "Serialization"]
uuid = "1a1011a3-84de-559e-8e89-a11a2f7dc383"

[[deps.SimpleDiffEq]]
deps = ["DiffEqBase", "LinearAlgebra", "MuladdMacro", "Parameters", "RecursiveArrayTools", "Reexport", "StaticArrays"]
git-tree-sha1 = "736e6abb40125e753c7ea407d3c15eba13b803ed"
uuid = "05bca326-078c-5bf0-a5bf-ce7c7982d7fd"
version = "1.8.0"

[[deps.SimpleNonlinearSolve]]
deps = ["ArrayInterfaceCore", "FiniteDiff", "ForwardDiff", "Reexport", "SciMLBase", "SnoopPrecompile", "StaticArraysCore"]
git-tree-sha1 = "fc4b9f81a033cf6879c91bb7f5b3ff59008c7dd2"
uuid = "727e6d20-b764-4bd8-a329-72de5adea6c7"
version = "0.1.4"

[[deps.SimpleTraits]]
deps = ["InteractiveUtils", "MacroTools"]
git-tree-sha1 = "5d7e3f4e11935503d3ecaf7186eac40602e7d231"
uuid = "699a6c99-e7fa-54fc-8d76-47d257e15c1d"
version = "0.9.4"

[[deps.Snappy]]
deps = ["CEnum", "snappy_jll"]
git-tree-sha1 = "72bae53c0691f4b6fd259587dab8821ae0e025f6"
uuid = "59d4ed8c-697a-5b28-a4c7-fe95c22820f9"
version = "0.4.2"

[[deps.SnoopPrecompile]]
deps = ["Preferences"]
git-tree-sha1 = "e760a70afdcd461cf01a575947738d359234665c"
uuid = "66db9d55-30c0-4569-8b51-7e840670fc0c"
version = "1.0.3"

[[deps.Sockets]]
uuid = "6462fe0b-24de-5631-8697-dd941f90decc"

[[deps.SoftGlobalScope]]
deps = ["REPL"]
git-tree-sha1 = "986ec2b6162ccb95de5892ed17832f95badf770c"
uuid = "b85f4697-e234-5449-a836-ec8e2f98b302"
version = "1.1.0"

[[deps.SortingAlgorithms]]
deps = ["DataStructures"]
git-tree-sha1 = "a4ada03f999bd01b3a25dcaa30b2d929fe537e00"
uuid = "a2af1166-a08f-5f64-846c-94a0d3cef48c"
version = "1.1.0"

[[deps.SparseArrays]]
deps = ["LinearAlgebra", "Random"]
uuid = "2f01184e-e22b-5df5-ae63-d93ebab69eaf"

[[deps.SparseDiffTools]]
deps = ["Adapt", "ArrayInterfaceCore", "ArrayInterfaceStaticArrays", "Compat", "DataStructures", "FiniteDiff", "ForwardDiff", "Graphs", "LinearAlgebra", "Requires", "SparseArrays", "StaticArrays", "VertexSafeGraphs"]
git-tree-sha1 = "4245283bee733122a9cb4545748d64e0c63337c0"
uuid = "47a9eef4-7e08-11e9-0b38-333d64bd3804"
version = "1.30.0"

[[deps.Sparspak]]
deps = ["Libdl", "LinearAlgebra", "Logging", "OffsetArrays", "Printf", "SparseArrays", "Test"]
git-tree-sha1 = "4149bb50ccf65bc9c9d55e7001ca7aa4a7649603"
uuid = "e56a9233-b9d6-4f03-8d0f-1825330902ac"
version = "0.3.4"

[[deps.SpecialFunctions]]
deps = ["ChainRulesCore", "IrrationalConstants", "LogExpFunctions", "OpenLibm_jll", "OpenSpecFun_jll"]
git-tree-sha1 = "d75bda01f8c31ebb72df80a46c88b25d1c79c56d"
uuid = "276daf66-3868-5448-9aa4-cd146d93841b"
version = "2.1.7"

[[deps.SplittablesBase]]
deps = ["Setfield", "Test"]
git-tree-sha1 = "e08a62abc517eb79667d0a29dc08a3b589516bb5"
uuid = "171d559e-b47b-412a-8079-5efa626c420e"
version = "0.1.15"

[[deps.Static]]
deps = ["IfElse"]
git-tree-sha1 = "c35b107b61e7f34fa3f124026f2a9be97dea9e1c"
uuid = "aedffcd0-7271-4cad-89d0-dc628f76c6d3"
version = "0.8.3"

[[deps.StaticArrays]]
deps = ["LinearAlgebra", "Random", "StaticArraysCore", "Statistics"]
git-tree-sha1 = "6954a456979f23d05085727adb17c4551c19ecd1"
uuid = "90137ffa-7385-5640-81b9-e52037218182"
version = "1.5.12"

[[deps.StaticArraysCore]]
git-tree-sha1 = "6b7ba252635a5eff6a0b0664a41ee140a1c9e72a"
uuid = "1e83bf80-4336-4d27-bf5d-d5a4f845583c"
version = "1.4.0"

[[deps.Statistics]]
deps = ["LinearAlgebra", "SparseArrays"]
uuid = "10745b16-79ce-11e8-11f9-7d13ad32a3b2"

[[deps.StatsAPI]]
deps = ["LinearAlgebra"]
git-tree-sha1 = "f9af7f195fb13589dd2e2d57fdb401717d2eb1f6"
uuid = "82ae8749-77ed-4fe6-ae5f-f523153014b0"
version = "1.5.0"

[[deps.StatsBase]]
deps = ["DataAPI", "DataStructures", "LinearAlgebra", "LogExpFunctions", "Missings", "Printf", "Random", "SortingAlgorithms", "SparseArrays", "Statistics", "StatsAPI"]
git-tree-sha1 = "d1bf48bfcc554a3761a133fe3a9bb01488e06916"
uuid = "2913bbd2-ae8a-5f71-8c99-4fb6c76f3a91"
version = "0.33.21"

[[deps.StatsFuns]]
deps = ["ChainRulesCore", "HypergeometricFunctions", "InverseFunctions", "IrrationalConstants", "LogExpFunctions", "Reexport", "Rmath", "SpecialFunctions"]
git-tree-sha1 = "ab6083f09b3e617e34a956b43e9d51b824206932"
uuid = "4c63d2b9-4356-54db-8cca-17b64c39e42c"
version = "1.1.1"

[[deps.SteadyStateDiffEq]]
deps = ["DiffEqBase", "DiffEqCallbacks", "LinearAlgebra", "NLsolve", "Reexport", "SciMLBase"]
git-tree-sha1 = "32d8ad240e065b5c86088c759670cb6bcc96e45a"
uuid = "9672c7b4-1e72-59bd-8a11-6ac3964bc41f"
version = "1.11.0"

[[deps.StochasticDiffEq]]
deps = ["Adapt", "ArrayInterface", "DataStructures", "DiffEqBase", "DiffEqNoiseProcess", "DocStringExtensions", "FillArrays", "FiniteDiff", "ForwardDiff", "JumpProcesses", "LevyArea", "LinearAlgebra", "Logging", "MuladdMacro", "NLsolve", "OrdinaryDiffEq", "Random", "RandomNumbers", "RecursiveArrayTools", "Reexport", "SciMLBase", "SparseArrays", "SparseDiffTools", "StaticArrays", "UnPack"]
git-tree-sha1 = "c35d43a21e91fd53ebe31bd6a3d81745e1c8fca0"
uuid = "789caeaf-c7a9-5a7d-9973-96adeb23e2a0"
version = "6.57.4"

[[deps.StrideArraysCore]]
deps = ["ArrayInterface", "CloseOpenIntervals", "IfElse", "LayoutPointers", "ManualMemory", "SIMDTypes", "Static", "ThreadingUtilities"]
git-tree-sha1 = "70b6ee0e5cc1745a28dd9ba040b8e5ee28fffc69"
uuid = "7792a7ef-975c-4747-a70f-980b88e8d1da"
version = "0.4.5"

[[deps.StringManipulation]]
git-tree-sha1 = "46da2434b41f41ac3594ee9816ce5541c6096123"
uuid = "892a3eda-7b42-436c-8928-eab12a02cf0e"
version = "0.3.0"

[[deps.StructTypes]]
deps = ["Dates", "UUIDs"]
git-tree-sha1 = "ca4bccb03acf9faaf4137a9abc1881ed1841aa70"
uuid = "856f2bd8-1eba-4b0a-8007-ebc267875bd4"
version = "1.10.0"

[[deps.SuiteSparse]]
deps = ["Libdl", "LinearAlgebra", "Serialization", "SparseArrays"]
uuid = "4607b0f0-06f3-5cda-b6b1-a6196a1729e9"

[[deps.SuiteSparse_jll]]
deps = ["Artifacts", "Libdl", "Pkg", "libblastrampoline_jll"]
uuid = "bea87d4a-7f5b-5778-9afe-8cc45184846c"
version = "5.10.1+0"

[[deps.Sundials]]
deps = ["CEnum", "DataStructures", "DiffEqBase", "Libdl", "LinearAlgebra", "Logging", "Reexport", "SciMLBase", "SnoopPrecompile", "SparseArrays", "Sundials_jll"]
git-tree-sha1 = "3f8c27118d25ac5cfd36ec382c9c3a834c4d91ad"
uuid = "c3572dad-4567-51f8-b174-8c6c989267f4"
version = "4.11.4"

[[deps.Sundials_jll]]
deps = ["Artifacts", "CompilerSupportLibraries_jll", "JLLWrappers", "Libdl", "OpenBLAS_jll", "Pkg", "SuiteSparse_jll"]
git-tree-sha1 = "04777432d74ec5bc91ca047c9e0e0fd7f81acdb6"
uuid = "fb77eaff-e24c-56d4-86b1-d163f2edb164"
version = "5.2.1+0"

[[deps.SymbolicIndexingInterface]]
deps = ["DocStringExtensions"]
git-tree-sha1 = "6b764c160547240d868be4e961a5037f47ad7379"
uuid = "2efcf032-c050-4f8e-a9bb-153293bab1f5"
version = "0.2.1"

[[deps.SymbolicUtils]]
deps = ["AbstractTrees", "Bijections", "ChainRulesCore", "Combinatorics", "ConstructionBase", "DataStructures", "DocStringExtensions", "DynamicPolynomials", "IfElse", "LabelledArrays", "LinearAlgebra", "Metatheory", "MultivariatePolynomials", "NaNMath", "Setfield", "SparseArrays", "SpecialFunctions", "StaticArrays", "TermInterface", "TimerOutputs"]
git-tree-sha1 = "027b43d312f6d52187bb16c2d4f0588ddb8c4bb2"
uuid = "d1185830-fcd6-423d-90d6-eec64667417b"
version = "0.19.11"

[[deps.Symbolics]]
deps = ["ArrayInterfaceCore", "ConstructionBase", "DataStructures", "DiffRules", "Distributions", "DocStringExtensions", "DomainSets", "Groebner", "IfElse", "LaTeXStrings", "LambertW", "Latexify", "Libdl", "LinearAlgebra", "MacroTools", "Markdown", "Metatheory", "NaNMath", "RecipesBase", "Reexport", "Requires", "RuntimeGeneratedFunctions", "SciMLBase", "Setfield", "SparseArrays", "SpecialFunctions", "StaticArrays", "SymbolicUtils", "TermInterface", "TreeViews"]
git-tree-sha1 = "111fbf43883d95989577133aeeb889f2040d0aea"
uuid = "0c5d862f-8b57-4792-8d23-62f2024744c7"
version = "4.14.0"

[[deps.TOML]]
deps = ["Dates"]
uuid = "fa267f1f-6049-4f14-aa54-33bafae1ed76"
version = "1.0.0"

[[deps.TableOperations]]
deps = ["SentinelArrays", "Tables", "Test"]
git-tree-sha1 = "e383c87cf2a1dc41fa30c093b2a19877c83e1bc1"
uuid = "ab02a1b2-a7df-11e8-156e-fb1833f50b87"
version = "1.2.0"

[[deps.TableTraits]]
deps = ["IteratorInterfaceExtensions"]
git-tree-sha1 = "c06b2f539df1c6efa794486abfb6ed2022561a39"
uuid = "3783bdb8-4a98-5b6b-af9a-565f29a5fe9c"
version = "1.0.1"

[[deps.Tables]]
deps = ["DataAPI", "DataValueInterfaces", "IteratorInterfaceExtensions", "LinearAlgebra", "OrderedCollections", "TableTraits", "Test"]
git-tree-sha1 = "1544b926975372da01227b382066ab70e574a3ec"
uuid = "bd369af6-aec1-5ad0-b16a-f7cc5008161c"
version = "1.10.1"

[[deps.Tar]]
deps = ["ArgTools", "SHA"]
uuid = "a4e569a6-e804-4fa4-b0f3-eef7a1d5b13e"
version = "1.10.1"

[[deps.TensorCore]]
deps = ["LinearAlgebra"]
git-tree-sha1 = "1feb45f88d133a655e001435632f019a9a1bcdb6"
uuid = "62fd8b95-f654-4bbd-a8a5-9c27f68ccd50"
version = "0.1.1"

[[deps.TermInterface]]
git-tree-sha1 = "7aa601f12708243987b88d1b453541a75e3d8c7a"
uuid = "8ea1fca8-c5ef-4a55-8b96-4e9afe9c9a3c"
version = "0.2.3"

[[deps.Test]]
deps = ["InteractiveUtils", "Logging", "Random", "Serialization"]
uuid = "8dfed614-e22c-5e08-85e1-65c5234f0b40"

[[deps.ThreadingUtilities]]
deps = ["ManualMemory"]
git-tree-sha1 = "f8629df51cab659d70d2e5618a430b4d3f37f2c3"
uuid = "8290d209-cae3-49c0-8002-c8c24d57dab5"
version = "0.5.0"

[[deps.ThreadsX]]
deps = ["ArgCheck", "BangBang", "ConstructionBase", "InitialValues", "MicroCollections", "Referenceables", "Setfield", "SplittablesBase", "Transducers"]
git-tree-sha1 = "34e6bcf36b9ed5d56489600cf9f3c16843fa2aa2"
uuid = "ac1d9e8a-700a-412c-b207-f0111f4b6c0d"
version = "0.1.11"

[[deps.Thrift]]
deps = ["CodecZlib", "CodecZstd", "Distributed", "Sockets", "ThriftJuliaCompiler_jll", "TranscodingStreams"]
git-tree-sha1 = "3da5858caabd351ee845bb1d547e4d70dbc65b20"
uuid = "8d9c9c80-f77e-5080-9541-c6f69d204e22"
version = "0.8.4"

[[deps.ThriftJuliaCompiler_jll]]
deps = ["Artifacts", "JLLWrappers", "Libdl", "Pkg"]
git-tree-sha1 = "949a51ca85d31b063531eed49e38a6c9b9bae58b"
uuid = "815b9798-8dd0-5549-95cc-3cf7d01bce66"
version = "0.12.1+0"

[[deps.TimeZones]]
deps = ["Dates", "Downloads", "InlineStrings", "LazyArtifacts", "Mocking", "Printf", "RecipesBase", "Scratch", "Unicode"]
git-tree-sha1 = "a92ec4466fc6e3dd704e2668b5e7f24add36d242"
uuid = "f269a46b-ccf7-5d73-abea-4c690281aa53"
version = "1.9.1"

[[deps.TimerOutputs]]
deps = ["ExprTools", "Printf"]
git-tree-sha1 = "f2fd3f288dfc6f507b0c3a2eb3bac009251e548b"
uuid = "a759f4b9-e2f1-59dc-863e-4aeb61b1ea8f"
version = "0.5.22"

[[deps.Tokenize]]
git-tree-sha1 = "90538bf898832b6ebd900fa40f223e695970e3a5"
uuid = "0796e94c-ce3b-5d07-9a54-7f471281c624"
version = "0.5.25"

[[deps.TranscodingStreams]]
deps = ["Random", "Test"]
git-tree-sha1 = "94f38103c984f89cf77c402f2a68dbd870f8165f"
uuid = "3bb67fe8-82b1-5028-8e26-92a6c54297fa"
version = "0.9.11"

[[deps.Transducers]]
deps = ["Adapt", "ArgCheck", "BangBang", "Baselet", "CompositionsBase", "DefineSingletons", "Distributed", "InitialValues", "Logging", "Markdown", "MicroCollections", "Requires", "Setfield", "SplittablesBase", "Tables"]
git-tree-sha1 = "c42fa452a60f022e9e087823b47e5a5f8adc53d5"
uuid = "28d57a85-8fef-5791-bfe6-a80928e7c999"
version = "0.4.75"

[[deps.TreeViews]]
deps = ["Test"]
git-tree-sha1 = "8d0d7a3fe2f30d6a7f833a5f19f7c7a5b396eae6"
uuid = "a2a6695c-b41b-5b7d-aed9-dbfdeacea5d7"
version = "0.3.0"

[[deps.TriangularSolve]]
deps = ["CloseOpenIntervals", "IfElse", "LayoutPointers", "LinearAlgebra", "LoopVectorization", "Polyester", "SnoopPrecompile", "Static", "VectorizationBase"]
git-tree-sha1 = "6cca884e0fe17916da63c62dc1bf5896ce5d723e"
uuid = "d5829a12-d9aa-46ab-831f-fb7c9ab06edf"
version = "0.1.17"

[[deps.Tricks]]
git-tree-sha1 = "6bac775f2d42a611cdfcd1fb217ee719630c4175"
uuid = "410a4b4d-49e4-4fbc-ab6d-cb71b17b3775"
version = "0.1.6"

[[deps.URIs]]
git-tree-sha1 = "ac00576f90d8a259f2c9d823e91d1de3fd44d348"
uuid = "5c2747f8-b7ea-4ff2-ba2e-563bfd36b1d4"
version = "1.4.1"

[[deps.UUIDs]]
deps = ["Random", "SHA"]
uuid = "cf7118a7-6976-5b1a-9a39-7adc72f591a4"

[[deps.UnPack]]
git-tree-sha1 = "387c1f73762231e86e0c9c5443ce3b4a0a9a0c2b"
uuid = "3a884ed6-31ef-47d7-9d2a-63182c4928ed"
version = "1.0.2"

[[deps.Unicode]]
uuid = "4ec0a83e-493e-50e2-b9ac-8f72acf5a8f5"

[[deps.Unitful]]
deps = ["ConstructionBase", "Dates", "LinearAlgebra", "Random"]
git-tree-sha1 = "d670a70dd3cdbe1c1186f2f17c9a68a7ec24838c"
uuid = "1986cc42-f94f-5a68-af5c-568840ba703d"
version = "1.12.2"

[[deps.UnsafeArrays]]
git-tree-sha1 = "3350f94f6caa02f324a23645bf524fc9334c7488"
uuid = "c4a57d5a-5b31-53a6-b365-19f8c011fbd6"
version = "1.0.4"

[[deps.UnsafeAtomics]]
git-tree-sha1 = "6331ac3440856ea1988316b46045303bef658278"
uuid = "013be700-e6cd-48c3-b4a1-df204f14c38f"
version = "0.2.1"

[[deps.UnsafeAtomicsLLVM]]
deps = ["LLVM", "UnsafeAtomics"]
git-tree-sha1 = "33af9d2031d0dc09e2be9a0d4beefec4466def8e"
uuid = "d80eeb9a-aca5-4d75-85e5-170c8b632249"
version = "0.1.0"

[[deps.VectorizationBase]]
deps = ["ArrayInterface", "CPUSummary", "HostCPUFeatures", "IfElse", "LayoutPointers", "Libdl", "LinearAlgebra", "SIMDTypes", "Static"]
git-tree-sha1 = "6b1dc4fc039d273abc247eba675ac1299380e5d9"
uuid = "3d5dd08c-fd9d-11e8-17fa-ed2836048c2f"
version = "0.21.57"

[[deps.VersionParsing]]
git-tree-sha1 = "58d6e80b4ee071f5efd07fda82cb9fbe17200868"
uuid = "81def892-9a0e-5fdd-b105-ffc91e053289"
version = "1.3.0"

[[deps.VertexSafeGraphs]]
deps = ["Graphs"]
git-tree-sha1 = "8351f8d73d7e880bfc042a8b6922684ebeafb35c"
uuid = "19fa3120-7c27-5ec5-8db8-b0b0aa330d6f"
version = "0.2.0"

[[deps.WeakRefStrings]]
deps = ["DataAPI", "InlineStrings", "Parsers"]
git-tree-sha1 = "b1be2855ed9ed8eac54e5caff2afcdb442d52c23"
uuid = "ea10d353-3f73-51f8-a26c-33c1cb351aa5"
version = "1.4.2"

[[deps.WoodburyMatrices]]
deps = ["LinearAlgebra", "SparseArrays"]
git-tree-sha1 = "de67fa59e33ad156a590055375a30b23c40299d3"
uuid = "efce3f68-66dc-5838-9240-27a6d6f5f9b6"
version = "0.5.5"

[[deps.WorkerUtilities]]
git-tree-sha1 = "cd1659ba0d57b71a464a29e64dbc67cfe83d54e7"
uuid = "76eceee3-57b5-4d4a-8e66-0e911cebbf60"
version = "1.6.1"

[[deps.ZMQ]]
deps = ["FileWatching", "Sockets", "ZeroMQ_jll"]
git-tree-sha1 = "356d2bdcc0bce90aabee1d1c0f6d6f301eda8f77"
uuid = "c2297ded-f4af-51ae-bb23-16f91089e4e1"
version = "1.2.2"

[[deps.ZeroMQ_jll]]
deps = ["Artifacts", "JLLWrappers", "Libdl", "Pkg", "libsodium_jll"]
git-tree-sha1 = "fe5c65a526f066fb3000da137d5785d9649a8a47"
uuid = "8f1865be-045e-5c20-9c9f-bfbfb0764568"
version = "4.3.4+0"

[[deps.Zlib_jll]]
deps = ["Libdl"]
uuid = "83775a58-1f1d-513f-b197-d71354ab007a"
version = "1.2.12+3"

[[deps.Zstd_jll]]
deps = ["Artifacts", "JLLWrappers", "Libdl", "Pkg"]
git-tree-sha1 = "e45044cd873ded54b6a5bac0eb5c971392cf1927"
uuid = "3161d3a3-bdf6-5164-811a-617609db77b4"
version = "1.5.2+0"

[[deps.ZygoteRules]]
deps = ["MacroTools"]
git-tree-sha1 = "8c1a8e4dfacb1fd631745552c8db35d0deb09ea0"
uuid = "700de1a5-db45-46bc-99cf-38207098b444"
version = "0.2.2"

[[deps.libblastrampoline_jll]]
deps = ["Artifacts", "Libdl", "OpenBLAS_jll"]
uuid = "8e850b90-86db-534c-a0d3-1478176c7d93"
version = "5.1.1+0"

[[deps.libsodium_jll]]
deps = ["Artifacts", "JLLWrappers", "Libdl", "Pkg"]
git-tree-sha1 = "848ab3d00fe39d6fbc2a8641048f8f272af1c51e"
uuid = "a9144af2-ca23-56d9-984f-0d03f7b5ccf8"
version = "1.0.20+0"

[[deps.nghttp2_jll]]
deps = ["Artifacts", "Libdl"]
uuid = "8e850ede-7688-5339-a07c-302acd2aaf8d"
version = "1.48.0+0"

[[deps.p7zip_jll]]
deps = ["Artifacts", "Libdl"]
uuid = "3f19e933-33d8-53b3-aaab-bd5110c3b7a0"
version = "17.4.0+0"

[[deps.snappy_jll]]
deps = ["Artifacts", "JLLWrappers", "LZO_jll", "Libdl", "Pkg", "Zlib_jll"]
git-tree-sha1 = "985c1da710b0e43f7c52f037441021dfd0e3be14"
uuid = "fe1e1685-f7be-5f59-ac9f-4ca204017dfd"
version = "1.1.9+1"
```
