## Supplementary material for "Examining the role of oxygen-binding proteins on the early evolution of multicellularity": Myoglobin diffusion model code: Project.toml

```
[deps]
Arrow = "69666777-d1a9-59fb-9406-91d4454c9d45"
BenchmarkTools = "6e4b80f9-dd63-53aa-95a3-0cdb28fa8baf"
CSV = "336ed68f-0bac-5ca0-87d4-7b16caf5d00b"
CUDA = "052768ef-5323-5732-b1bb-66c8b64840ba"
ColorSchemes = "35d6a980-a343-548e-a6ea-1d62b119f2f4"
DataFrames = "a93c6f00-e57d-5684-b7b6-d8193f3e46c0"
DiffEqGPU = "071ae1c0-96b5-11e9-1965-c90190d839ea"
DifferentialEquations = "0c46a032-eb83-5123-abaf-570d42b7fbaa"
DomainSets = "5b8099bc-c8ec-5219-889f-1d9e522a28bf"
IJulia = "7073ff75-c697-5162-941a-fcdaad2a7d2a"
Interpolations = "a98d9a8b-a2ab-59e6-89dd-64a1c18fca59"
MethodOfLines = "94925ecb-adb7-4558-8ed8-f975c56a0bf4"
ModelingToolkit = "961ee093-0014-501f-94e3-6117800e7a78"
NonlinearSolve = "8913a72c-1f9b-4ce2-8d82-65094dcecaec"
Parquet2 = "98572fba-bba0-415d-956f-fa77e587d26d"
Pkg = "44cfe95a-1eb2-52ea-b672-e2afdf69b78f"
StaticArrays = "90137ffa-7385-5640-81b9-e52037218182"
Tables = "bd369af6-aec1-5ad0-b16a-f7cc5008161c"
```
